# Feed-forward loops by NR5A2 ensure robust gene activation during pre-implantation development

**DOI:** 10.1101/2025.02.14.638292

**Authors:** Wataru Kobayashi, Siwat Ruangroengkulrith, Eda Nur Arslantas, Adarsh Mohanan, Kikuë Tachibana

## Abstract

The pioneer transcription factor NR5A2 plays multiple roles in regulating zygotic genome activation and expression of lineage-determining factor during mouse pre-implantation development. However, how NR5A2 differentially regulates transcriptional networks at distinct developmental stages remains unknown. Here, we demonstrate the dynamics of chromatin binding profiles of NR5A2 from the 2-cell to morula stage, which corresponds to the totipotency-to-pluripotency transition. Our NR5A2 CUT&Tag analysis identifies lineage-determining factor KLF and GATA families as co-regulators for NR5A2 in mouse embryos. We find that KLF5 cooperates with NR5A2 to enhance H3K27ac deposition, and NR5A2 predominantly promotes chromatin accessibility. NR5A2 regulates *Xist* expression either directly or indirectly through its role in up-regulating GATA factor expression. *In vitro* assays reveal that NR5A2 co-binds to nucleosomes with KLF5 and GATA6, suggesting that these pioneer transcription factors can simultaneously bind to the chromatin. Our findings highlight the role of NR5A2 in ensuring robust gene activation during pre-implantation development through feed-forward regulatory loops with lineage-determining transcription factors.

## Introduction

Upon fertilization, the zygote acquires totipotency, which is the developmental potential to generate all cell types and form a complete organism. In mice, totipotency is maintained through zygote and 2-cell stages (Tarkowski, 1959), and the potential of totipotency gradually decreases during cleavage divisions until reaching a pluripotent or differentiated state (Kelly, 1977; Rossant, 1976). Initially, transcriptionally silent embryos are awakened by transcription referred to as zygotic genome activation (ZGA) (Jukam *et al*, 2017; Kravchenko & Tachibana, 2025). Following ZGA, embryos undergo a series of cleavage-stage divisions and develop into blastocysts consisting of inner cell mass (ICM) and trophectoderm (TE) (Rossant, 2018; Rossant & Tam, 2009; Zernicka-Goetz *et al*, 2009). The ICM subsequently gives rise to pluripotent epiblast (Epi) and primitive endoderm (PrE) (Rossant, 2018; Rossant & Tam, 2009; Zernicka-Goetz *et al*., 2009). These developmental transitions are driven by dynamic transcriptional regulatory networks that govern cell fate specification during pre-implantation development.

Pioneer transcription factors play critical roles in facilitating chromatin remodeling and driving cell fate transitions (Barral & Zaret, 2024; Iwafuchi-Doi & Zaret, 2014; Zaret & Carroll, 2011). Among these, Krüppel-like factors (KLF) and GATA transcription factors (TFs) are known as regulators of early embryonic development. The KLF and GATA families consist of 17 and 6 members, respectively, with both overlapping and distinct roles. For example, KLF17, maternal inherited factor, is required for ZGA at the 2-cell stage (Hu *et al*, 2024), whereas KLF4 and KLF5 are expressed during ZGA and contribute in lineage specification towards ICM and TE (Ema *et al*, 2008; Kinisu *et al*, 2021; Lin *et al*, 2010). GATA4 and GATA6 are known as PrE lineage markers (Koutsourakis *et al*, 1999; Morrisey *et al*, 1998), while GATA3 contributes to TE lineage specification (Gerri *et al*, 2020; Home *et al*, 2009; Ralston *et al*, 2010). Highly conserved DNA binding domain conservation within these families suggest both functional redundancy and cell stage-specific functions. However, the detailed regulatory mechanisms underlying pioneer factor functions in mammalian pre-implantation remains poorly understood.

The orphan nuclear receptor NR5A2 is a critical pioneer transcription factor essential for pre-implantation development in mice (Festuccia *et al*, 2024; Gassler *et al*, 2022; Lai *et al*, 2023; Zhao *et al*, 2024). NR5A2 regulates ZGA in 2-cell stage (Gassler *et al*., 2022) and controls the expression of lineage-determining factors at the 8-cell stage (Festuccia *et al*., 2024; Lai *et al*., 2023; Zhao *et al*., 2024). NR5A2 is expressed in mouse embryonic stem cells (mESCs) and maintains naïve pluripotency (Festuccia *et al*, 2021). Mechanistically, NR5A2 facilitates gene expression by unwrapping nucleosomal DNA through minor groove anchor competition (Kobayashi *et al*, 2024). Thus, NR5A2 orchestrates distinct transcriptional regulatory networks during the totipotency-to-pluripotency transition. Our previous findings indicate that NR5A2 targets cell-type specific enhancer-like regions between 2-cell embryos and mESCs (Gassler *et al*., 2022). However, the mechanisms by which NR5A2 differentially regulates transcriptional networks at distinct developmental stages remain unknown (Li *et al*, 2024). Given that pioneer TFs bind to chromatin in a context-dependent manner, we hypothesized that NR5A2 functions with lineage-determining TFs to control distinct transcriptional networks during pre-implantation development.

## Results

### The dynamics of NR5A2 binding during the totipotency-to-pluripotency transition

To investigate the transcriptional networks regulated by NR5A2, we first determined its chromatin binding profiles during the totipotency-to-pluripotency transition. Using CUT&Tag of approximately 800 cells per stage (Kaya-Okur *et al*, 2019), genome-wide NR5A2 binding profiles were determined from the 2-cell to the morula stage (Fig. 1A and EV1B). As a reference, previously published ChIP-seq data (Atlasi *et al*, 2019) in embryonic stem cells (ESCs) are shown. *De novo* motif analysis showed that the peaks are enriched for the NR5A2 motif in each cell stage (Fig. 1B). Our analysis detected 5,836, 9,251, 19,839, and 14,854 reproducible peaks in 2-cell, 4-cell, 8-cell, and morula, respectively (Table S1). This data suggests that the NR5A2 binding peaks at the 8-cell stage and decreases from the morula stage. Alluvial plots showed a gain and loss of NR5A2 peaks during cell stage transitions. Although we cannot exclude that non-peaks could be a false negative due to the small cell numbers, the overall picture emerges that NR5A2 dynamically changes its chromatin binding with each cell cycle (Fig. 1C).

**Figure 1.**
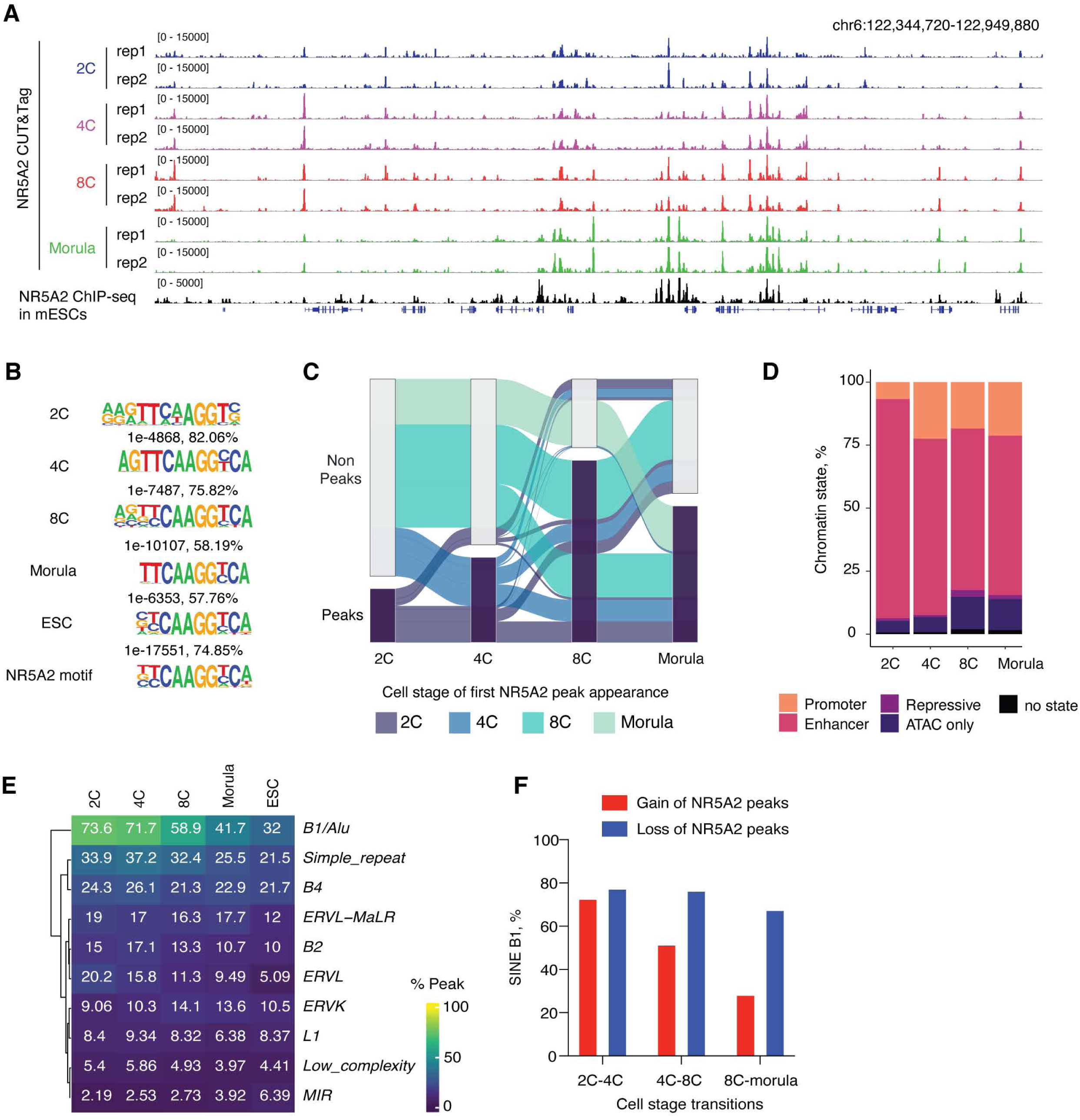
NR5A2 chromatin binding dynamics from the 2-cell to morula stage. (**A**) Integrated Genome Viewer (IGV) snapshot showing NR5A2 CUT&Tag signal across developmental stages: 2-cell (blue), 4-cell (purple), 8-cell (red), and morula (green), with two biological replicates. NR5A2 ChIP-seq data in mouse embryonic stem cells (mESCs, black) is shown for reference. (**B**) DNA motifs identified by HOMER *de novo* motif analysis of NR5A2 peaks at each developmental stage. The NR5A2 motif (MA0505.1) from the JASPAR database is shown as a reference. P-values and percentage of motif presence in peaks are shown. (**C**) Alluvial plots illustrating the dynamic changes in NR5A2 chromatin binding in each cell stage transition. (**D**) Classification of chromatin states associated with NR5A2 binding at each cell stage. Chromatin states were identified using ChromHMM. (**E**) Heatmaps depicting the enrichment of repetitive elements (subfamilies) within NR5A2 peaks (**F**) Bar chart representing the gain (red) and loss (blue) of NR5A2 peaks in each developmental transition.

NR5A2 predominantly targets putative enhancer regions including retrotransposon element *SINE B1*/*Alu* in the 2-cell stage when ZGA occurs (Gassler *et al*., 2022; Kravchenko & Tachibana, 2025). To investigate chromatin states targeted by NR5A2 at each cell stage, we applied hidden Markov model (HMM) to infer putative *cis*-regulatory elements (cREs) from chromatin accessibility and histone modifications using ChromHMM (Ernst & Kellis, 2017). To this end, H3K27ac CUT&Tag and omni ATAC-seq were conducted, and the profiles of H3K4me3, H3K9me3, and H3K27me3 were obtained from public data sets (Fig. EV1A, C-E and Table S2). Our analysis showed that NR5A2 preferentially targets promoters and putative enhancers from 2C to morula stages, which implies that NR5A2 is required for transcriptional activation (Fig. 1D). Consist with our previous report (Gassler *et al*., 2022), 73.6% NR5A2 peaks overlapped with *SINEB1/Alu* at the 2-cell stage. Intriguingly, the proportion of *SINE B1*/*Alu* in NR5A2 peaks decreased gradually from the 8-cell stage, dropping to 42% in the morula and 32% in ESCs (Fig. 1E). NR5A2 peaks at *SINE B1/Alu* were strongly gained at the 2-cell-to-4-cell transition and gradually decreased in the 8-cell-to-morula transition (Fig. 1F). In contrast, the loss of NR5A2 peaks remained relatively constant with each cell stage transition (Fig. 1F). These results suggest that NR5A2 specifically targets the retrotransposon elements *SINE B1/Alu* until cleavage-stage blastomeres, which correspond to the intermediate of the totipotency-pluripotency transition.

### KLF5 and GATA6 with NR5A2 co-occupy active chromatin

To identify potential co-regulators for NR5A2 in different cell stages, we investigated TF motifs enriched in NR5A2 peaks. The OCT4-SOX2 motif was highly enriched in mESC (Fig. 2A), suggesting that NR5A2 functions with OCT4 and SOX2 in pluripotent cells (Festuccia *et al*., 2021). However, the OCT4-SOX2 motif was not enriched in NR5A2 peaks from 2-cell to morula stage. Instead, KLF and GATA motifs in NR5A2 peaks are highly enriched at the 8-cell and morula stages (Fig. 2A). We then checked public Ribo-seq dataset (Xiong *et al*, 2022) and found that KLF5 and GATA6 are upregulated from the 8-cell to blastocyst stage among these family members (Fig. 2B). KLF5 are GATA6 are known as lineage-determining factors in mouse pre-implantation development. KLF5 is required to establish bi-potential cell fate for both ICM and TE gene expression (Ema *et al*., 2008; Kinisu *et al*., 2021; Lin *et al*., 2010), while GATA6 plays a critical role in PrE differentiation at the blastocyst stage (Koutsourakis *et al*., 1999; Morrisey *et al*., 1998; Wamaitha *et al*, 2015). These results indicate that NR5A2 may function with lineage-determining factors after ZGA.

**Figure 2.**
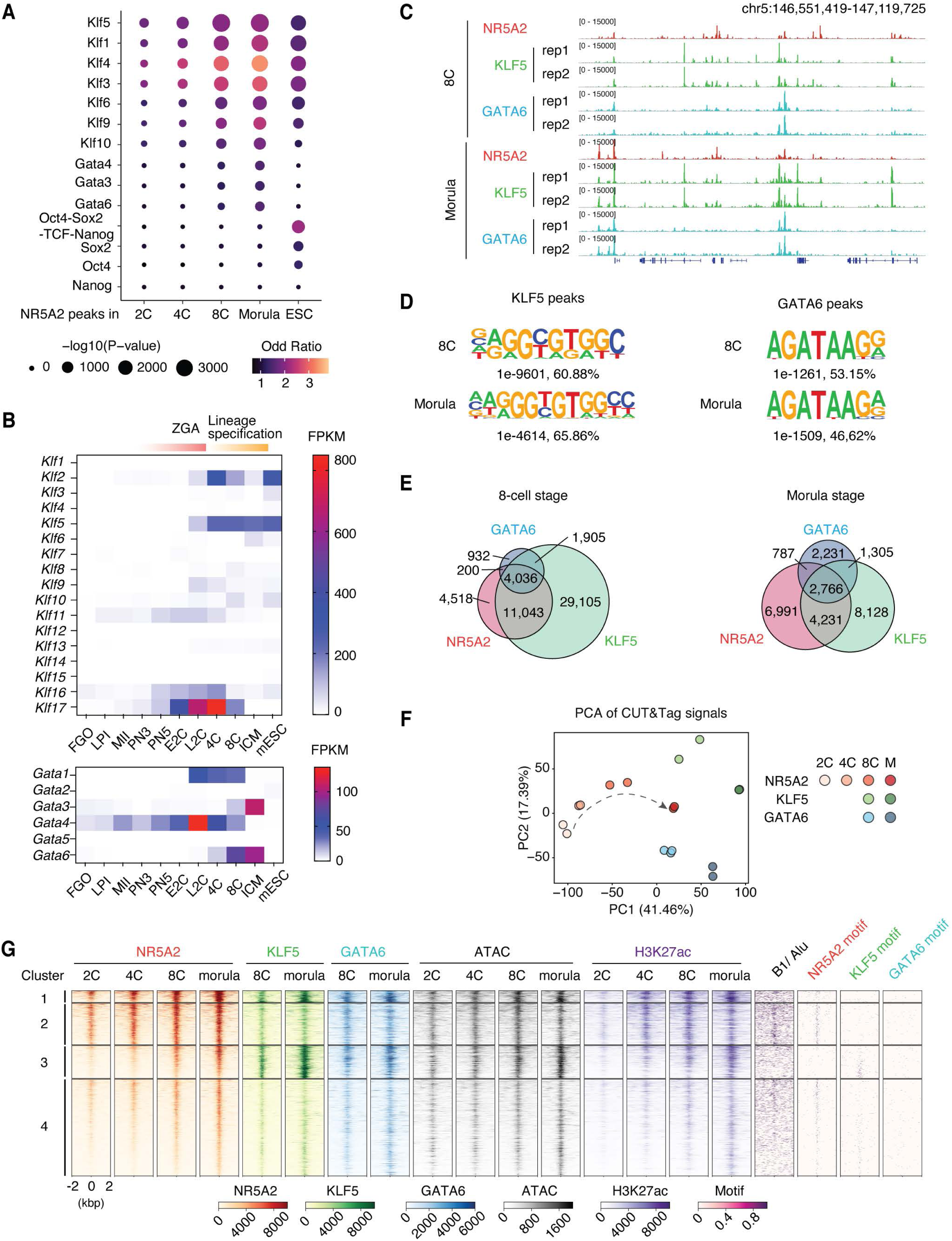
Identification of co-regulators for NR5A2. (**A**) Transcription factor motifs identified from NR5A2 peaks in each cell stage. The size of circle indicates p-value. Odd ratio indicates motif enrichment. (B) Ribo-seq data showing the expression level of KLF and GATA family. FGO: full-grown oocytes; LPI: Late prometaphase I oocytes; MII: Metaphase II oocytes; PN: Pronuclear stage; E2C: Early 2-cell embryo; L2C: Late 2-cell embryo; ICM: Inner cell mass. (**C**) IGV snapshot showing co-occupancy of KLF5 (green) and GATA6 (blue) with NR5A2 (red). CUT&Tag profiles for KLF5 and GATA6 at the 8-cell and morula stage are shown. (**D**) Venn diagrams showing the numbers of overlapped peaks between NR5A2, KLF5, and GATA6. (**E**) Heatmaps showing the enrichment of NR5A2, KLF5, GATA6, ATAC, H3K27ac, and transcription factor motif occurrence.

To test whether NR5A2 co-occupies chromatin with these lineage-determining factors, we performed KLF5 and GATA6 CUT&Tag on 8-cell and morula stages. As we expected, KLF5 and GATA6 peaks were detected at a subset of NR5A2 peaks, suggesting that these transcription factors co-occupy chromatin at some sites (Fig. 2C). *De novo* motif analysis confirmed the presence of canonical KLF5 and GATA motifs within peaks (Fig. 2D). The peaks of these three TFs overlapped at a subset of binding sites (Fig. 2E). Principal component analysis showed the trajectory of NR5A2 chromatin binding that follow the order of development. Interestingly, NR5A2 binding profile at morula stage tends to be similar to those of KLF5 and GATA6 at the 8-cell stage (Fig. 2F). To determine whether these co-occupied sites correlate with NR5A2 function at a later development stage, we classified NR5A2 peaks at the morula stage into 4 clusters based on signal of the three factors. We found a subset of Nr5a2 binding sites at morula stage that co-occupancy by all the three transcription factors. The co-binding of these factors correlated with the gain of H3K27ac levels at the morula stage (Fig. 2G, clusters 1 and 3). Consist with previous result, the newly gain NR5A2 binding sites preferentially target non-*SINE B1*/*Alu* regions with somewhat degenerated Nr5a2 motif (Fig. 2G, cluster 3 and EV1F). These data suggest that the potential co-occupancy of these three TFs is associated with the establishment of promoters or enhancers.

### NR5A2 and KLF5 function synergistically in embryonic development

To investigate whether NR5A2 regulates gene expression alongside KLF5 or GATA6 during early embryonic development, we performed siRNA-mediated knockdown to perturb their functions. We found that *Gata6* knockdown was inefficient up to the 8-cell stage (Fig. EV2A). Therefore, we focused on the cooperative role of KLF5 with NR5A2. To deplete both NR5A2 and KLF5 proteins, siRNAs targeting *Nr5a2* and *Klf5* were microinjected into zygotes and cultured until the 8-cell stage (Fig. 3A). Immunofluorescent staining of the embryos showed that knockdown of either *Nr5a2* or *Klf5* led to efficient depletion of the respective protein at the 8-cell stage (Fig. 3B and 3C). KLF5 levels were reduced by approximately 50% upon *Nr5a2* KD, suggesting that *Klf5* expression is regulated by NR5A2 (Fig. 3B and 3C). This is consistent with our previous finding that NR5A2 inhibition causes a significant downregulation of *Klf5* transcription in 2-cell embryos (Gassler et al., 2022). Double siRNA injection (*Nr5a2 Klf5* double KD) efficiently depleted both NR5A2 and KLF5 proteins (Fig. 3B and 3C).

**Figure 3.**
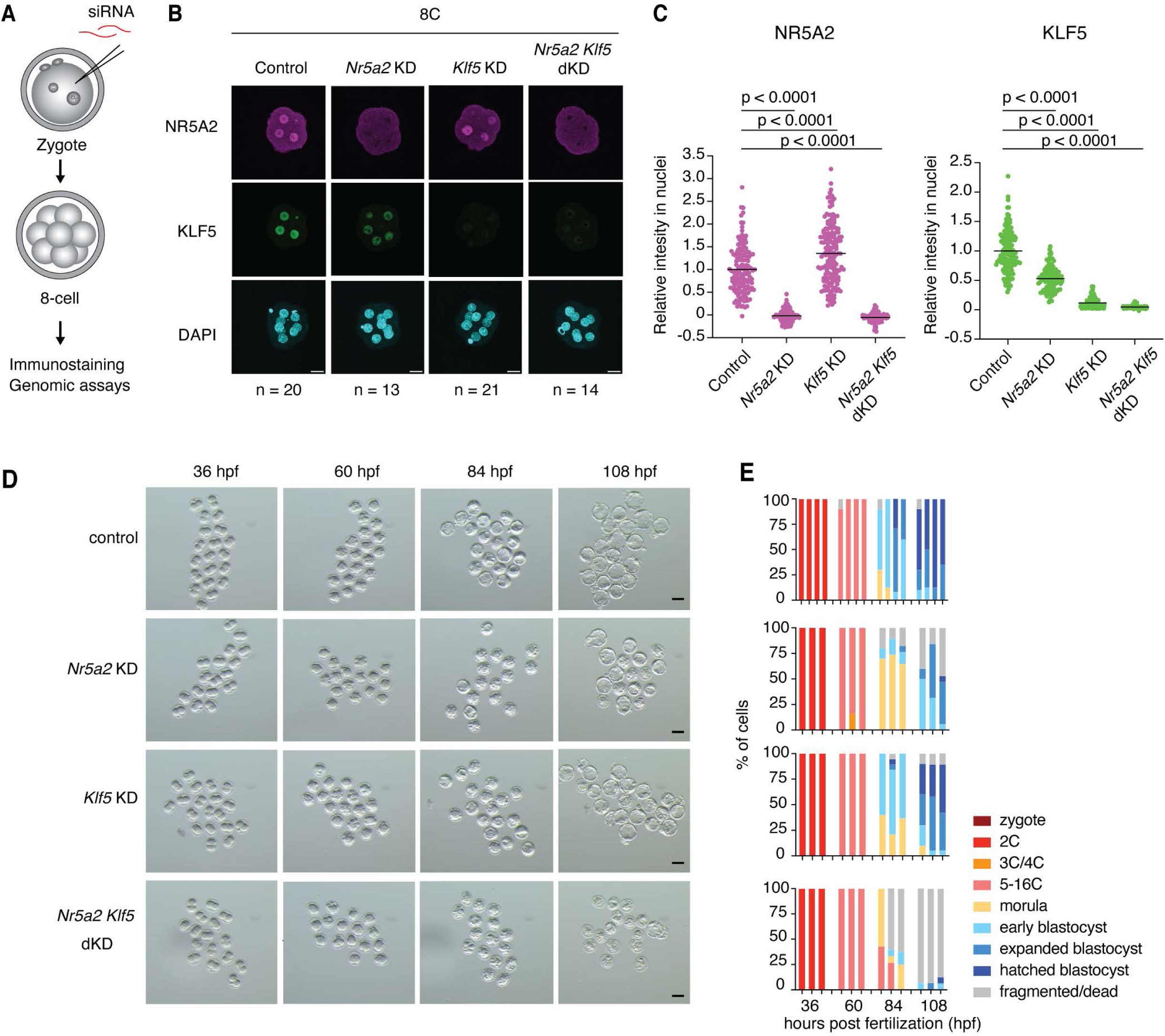
NR5A2 and KLF5 contribution to pre-implantation development. (**A**) Schematic representation of siRNA-mediated knockdown. siRNA targeting Nr5a2 and/or Klf5 was microinjected into zygotes. Immunostaining was then performed with 8-cell embryos (**B, C**) Representative images (**B**) and quantifications (**C**) of immunostaining analysis showing NR5A2 (magenta), KLF5 (green), and DAPI (cyan) of in 8-cell embryos. NR5A2 and KLF5 signals are shown in mid-section images. DAPI signals are presented as full z-stuck images to visualize all nuclei. The number of embryos examined (n) from three independent experiments is indicated. Scale bars, 20 μm. Bars overlaid on the plots indicate means. P values (t-test, two-sided) are shown. (**D**) Stereomicroscopic representative images showing embryonic development at different time points under different KD conditions. Scale bars, 75 μm. (**E**) Quantification of experiments of embryonic development. Sample sizes are as follows: control, n = 10, 8, 24, and 20 cells; *Nr5a2* KD, n = 10, 19, and 17; *Klf5* KD, n = 10, 19, and 19; *Nr5a2 Klf5* dKD, n = 14, 15, and 16.

Next, we tested whether NR5A2 and KLF5, alone or in combination, are required for development. Under our culture conditions, control embryos developed to the blastocyst stage, with more than 60% successfully hatching at 108 hpf (Fig. 3D and 3E). *Nr5a2* KD embryos developed with a delay and fragmented around the morula stage, consistent with previous work (Fig. 3D and 3E) (Festuccia *et al*., 2024; Lai *et al*., 2023; Zhao *et al*., 2024). In contrast, *Klf5* KD resulted in a mild developmental defect and blastocysts with a slightly smaller blastocoel (Fig. 3D and 3E). Notably, *Nr5a2 Klf5* double KD caused a severe developmental failure, with >90% of embryos dying and fragmenting (Fig. 3D and 3E). These results suggest that NR5A2 and KLF5 function synergistically in embryonic development.

### KlF5 cooperates with NR5A2 to enhance H3K27ac deposition for transcriptional activation

To investigate genome-wide transcriptional changes regulated by NR5A2 and KLF5, we performed KD experiments followed by RNA sequencing (RNA-seq) (Fig.3A, EV2B,C, and Table S3). *Nr5a2* KD and *Nr5a2 Klf5* dKD resulted in the down-regulation of 485 and 491 genes, respectively, with 354 genes commonly affected in both conditions (Fig. 4A and 4B). On the other hands, *Klf5* KD resulted in downregulation of merely 29 genes (Fig. 4A and 4B, discussed later), which could be due to inefficient knockdown (Fig. EV2B). These results suggest that NR5A2 regulates 72% (353 out of 491) of the genes down-regulated in *Nr5a2 Klf5* dKD, while an additional 128 genes were specifically down-regulated in *Nr5a2 Klf5* dKD. This raises the possibility that these 128 genes are potentially co-regulated by NR5A2 and KLF5.

**Figure 4.**
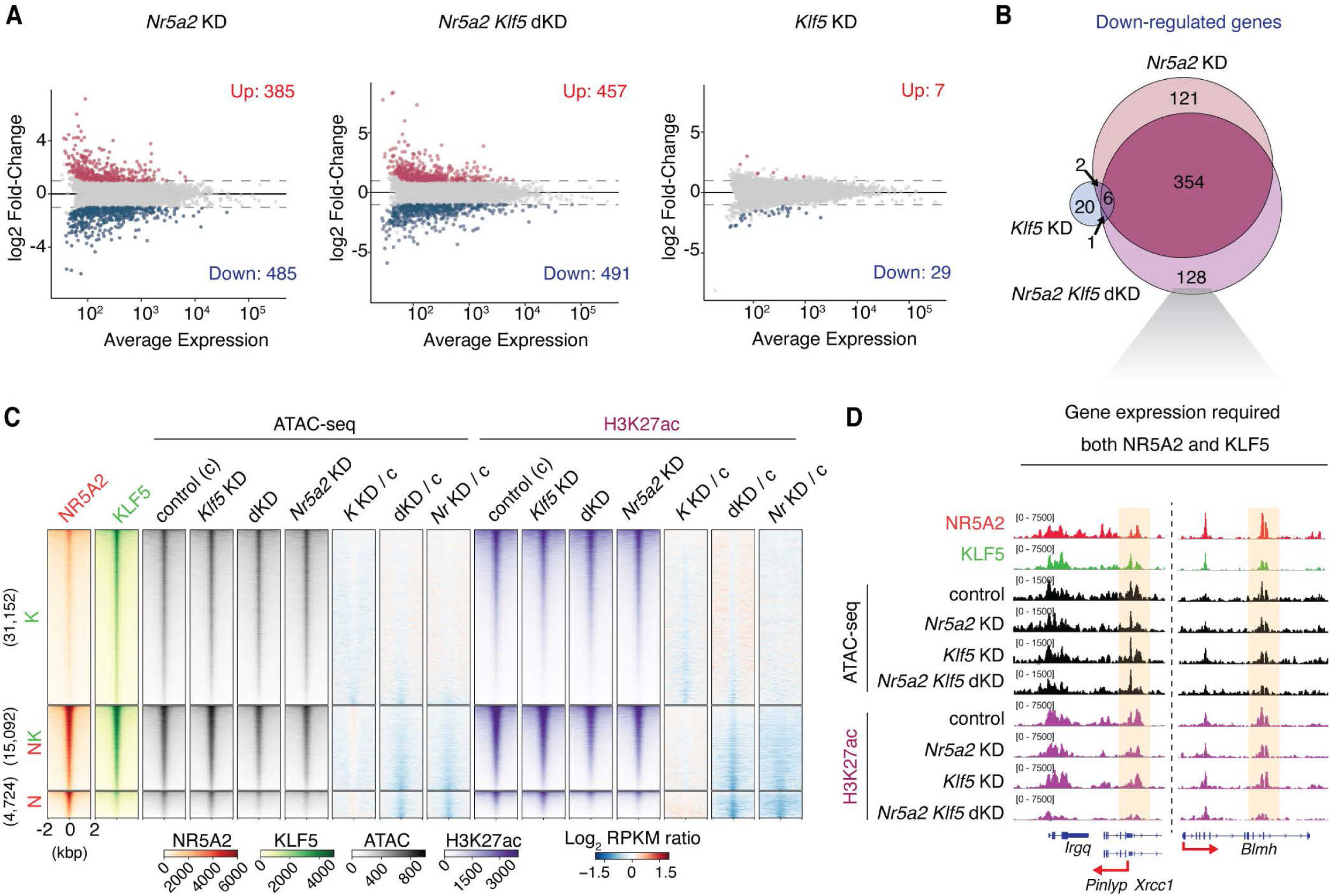
Transcriptional regulation by NR5A2 and KLF5 at the 8-cell stage. (**A**) MA plots of the log_2_ fold-change (*Nr5a2* KD/control, *Nr5a2 Klf5* dKD/control, and *Klf5* KD/control) in gene expression in *Nr5a2* KD (left), *Nr5a2 Klf5* dKD (center), and *Klf5* KD (right). (**B**) Venn diagrams showing the number of down-regulated genes overlapped in *Nr5a2* KD, *Nr5a2 Klf5* dKD, and *Klf5* KD. (**C**) Heatmaps showing enrichment of ATAC and H3K27ac on K, NK, and N regions. Log_2_ RPKM ratios (*Nr5a2* KD/control, *Nr5a2 Klf5* dKD/control, and *Klf5* KD/control) are shown, with data from two or three biological replicates merged. (**D**) IGV snapshot highlighting the representative genes regulated by both NR5A2 and KLF5. ATAC-seq and H3K27ac CUT&Tag are shown as merged profiles.

To elucidate the mechanism by which NR5A2 and KLF5 regulates chromatin accessibility and/or promoter/enhancer activity, we performed ATAC-seq (Corces *et al*, 2017) and H3K27ac CUT&Tag in each KD condition (Fig. 3A and EV2D, E). Since the efficiency of siRNA-mediated KD by microinjection can vary between experiments, we used the average signals of two to three replicates to avoid bias. To examine each factor contribution, NR5A2 and KLF5 binding at the 8-cell stage was classified into three categories: NR5A2-alone regions (N), KLF5-alone regions (K) and NR5A2 KLF5 co-occupied regions (NK). *Nr5a2* KD resulted in a decrease of chromatin accessibility at NR5A2-bound regions in agreement with previous reports (Festuccia *et al*., 2024; Lai *et al*., 2023). Additionally, H3K27ac levels were substantially decreased upon *Nr5a2* KD (Fig. 4C). IGV snapshots showed reductions in both chromatin accessibility and H3K27ac at regulatory elements of early ICM and TE genes (Fig. EV2F). These data are consistent with NR5A2’s function as a pioneer transcription factor that promotes chromatin accessibility and suggests an involvement in the deposition of H3K27ac at regulatory elements.

We then examined how NR5A2 and KLF5 promote transcriptional activation. *Klf5* KD had negligible effects on chromatin accessibility, but H3K27ac was mildly reduced at KLF5-bound regions (Fig. 4C). At NR5A2 KLF5 co-occupied regions, the reduction in chromatin accessibility upon *Nr5a2 Klf5* dKD was comparable to that observed in *Nr5a2* KD, suggesting that NR5A2 is primarily responsible for chromatin opening (Fig. 4C). Another possibility is that KLF5 depletion is stronger in the dKD because both its transcription (by NR5A2) is reduced and residual transcripts are knocked down. It therefore cannot be excluded that KLF5 contributes to chromatin opening. Interestingly, H3K27ac was further reduced in *Nr5a2 Klf5* dKD compared to *Nr5a2* KD (Fig. 4C). These data indicate that, while KLF5 may not directly influence chromatin accessibility, it contributes to H3K27ac deposition together with NR5A2 at co-occupied regions. Indeed, we observed a prominent decrease of H3K27ac at genes specifically down-regulated in *Nr5a2 Klf5* dKD (Fig. 4D). Taken together, these findings demonstrate that KLF5 cooperates with NR5A2 to enhance H3K27ac deposition for transcriptional activation.

### KLF5 directly regulates other KLF family members

The perturbation of transcription factors will affect not only their direct targets but potentially also other transcription factors and their targets. We therefore examined which other transcription factors are affected by *Klf5* KD. The 29 down-regulated genes in *Klf5* KD embryos (Fig. 4A and EV3A) include the pluripotency regulator *Prdm14* as well as several KLF family members, including *Klf3, Klf10*, and *Klf11* (Fig. EV3B). These genes are less down-regulated in *Nr5a2 Klf5* dKD compared to *Klf5* KD due to unknown reasons. The siRNA sequence targeting *Klf5* shows minimal overlap with other KLF family members, including NR5A2-regulated *Klf4*, suggesting that the effects are not due to off-targets (Fig. E3C). Furthermore, KLF5 CUT&Tag showed a strong enrichment at the promoters of these genes (Fig. EV3D), consistent with KLF5 directly regulating expression of other KLF family members. Previous studies showed that homozygous knockout mice for *Klf3^−/−^* (Sue *et al*, 2008), *Klf10^−/−^* (Yoo *et al*, 2023) and *Klf11^−/−^* (Song *et al*, 2005) are viable, suggesting that these KLF family members are possibly redundant. Interestingly, KLF3 protein is highly expressed in 8-cell stage and activates 8-cell activated genes in mESCs (Hao *et al*, 2020). This suggests that multiple KLF family members may be required to ensure proper development.

### NR5A2 regulates GATA factors and *Xist*

In early embryonic development, GATA transcription factors regulate the expression of *Xist,* a non-coding RNA essential for initiating X chromosome inactivation (XCI) (Ravid Lustig *et al*, 2023). Our RNA-seq data revealed that *Nr5a2* KD resulted in down-regulation of *Xist* and its regulator *Gata1* (Fig. 5A). While *Gata4* and *Gata6* showed only mild transcriptional down-regulation, immunofluorescence analysis showed a significant reduction in GATA6 protein upon *Nr5a2* KD (Fig. 5B and 5C). To rule out the possibility that our observed effect on XCI genes resulted from the stochastic imbalance of the ratio between male and female embryos in bulk RNA-seq, we re-analyzed published single-embryo RNA-seq data (Festuccia *et al*., 2024) with *Nr5a2* maternal-zygotic knockout 8-cell embryos. We found that *Xist* is substantially decreased in female embryos (Fig. EV4). These findings raise the interesting possibility that NR5A2 functions upstream of GATA transcription factors in regulating imprinted XCI.

**Figure 5.**
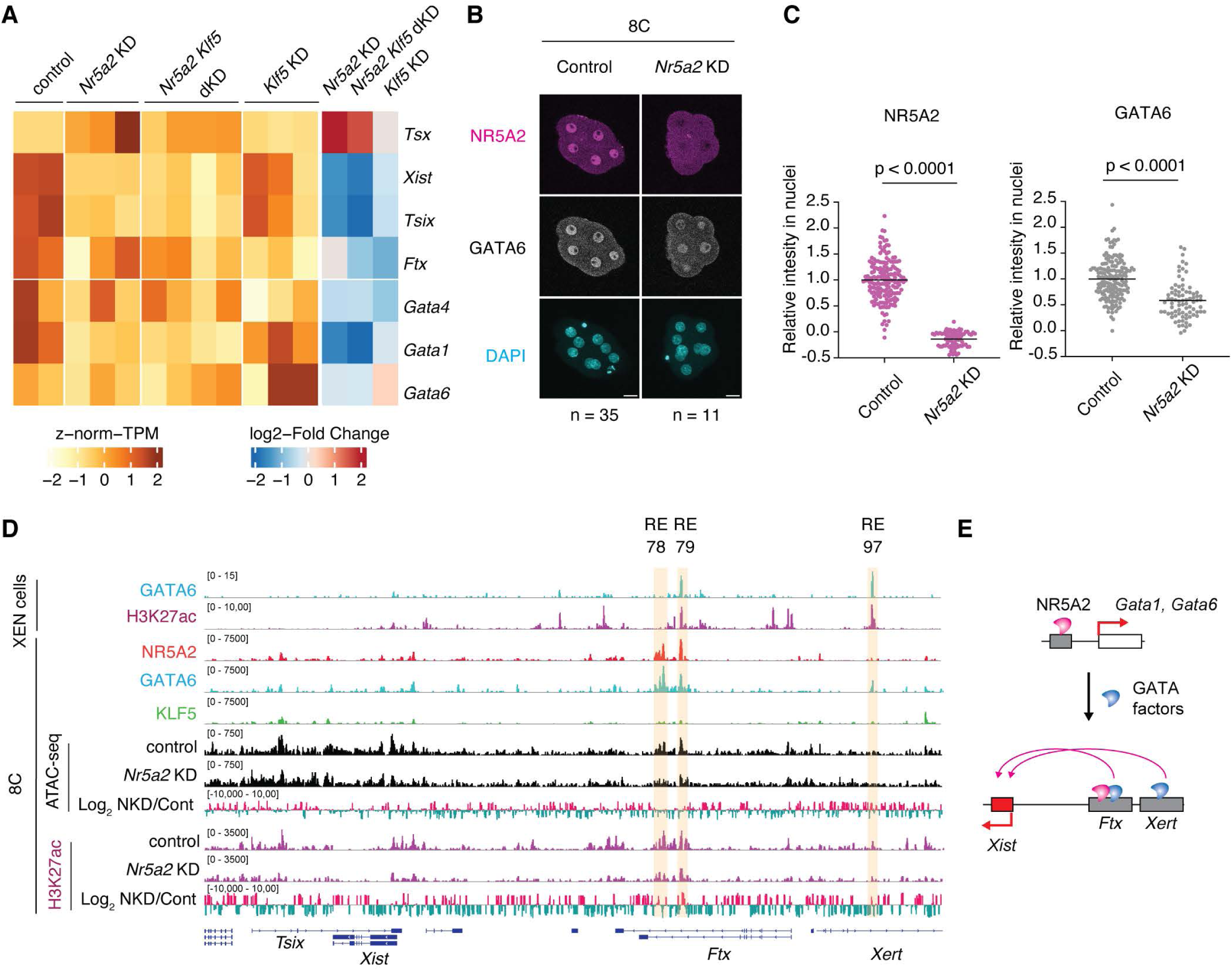
*Xist* expression is regulated by NR5A2 and GATA factors. (**A**) RNA expression of XCI-related genes and GATA family members. Heatmaps showing z-normalized TPM values for each replicate and log_2_ fold-changes (*Nr5a2* KD/control, *Nr5a2 Klf5* dKD/control, and *Klf5* KD/control). (**B, C**) Representative images (**B**) and quantifications (**C**) of immunostaining analysis showing NR5A2 (magenta), GATA6 (grey), and DAPI (cyan) in 8-cell embryos. NR5A2 and GATA signals are shown in mid-section images. DAPI signals are shown as full z-stuck images. The numbers of embryos examined (n) from three independent experiments are indicated. Scale bars, 20 μm. Bars overlaid on the plots indicate means. P values (t-test, two-sided) are shown. (**D**) IGV snapshot showing NR5A2 (red), GATA6 (blue), KLF5 (green) near *Xist* genome locus. GATA6 and H3K27ac signals from extraembryonic endoderm stem (XEN) cells are shown as control. Log_2_ fold-changes (*Nr5a2* KD/control) in ATAC-seq and H3K27ac are shown. (**E**) A model of *Xist* regulation by NR5A2 and GATA factors.

Previous studies identified distal regulatory elements (RE) 79 and 97 as *Xist* enhancers targeted by GATA transcription factors (Ravid Lustig *et al*., 2023). In our CUT&Tag data, GATA6 occupies RE79 and RE97 at the 8-cell stage (Fig. 5D), supporting a role of GATA6, and potentially other GATA family members, in targeting these distal REs to regulate imprinted XCI in pre-implantation development. Interestingly, we found that NR5A2 co-occupies RE79 with GATA6 but is not detected at RE97. Despite NR5A2 binding at RE79, chromatin accessibility and H3K27ac remained unchanged upon *Nr5a2* KD, which may indicate that NR5A2 binding is facilitated by residual GATA-dependent pioneering activity.

In addition to these REs, NR5A2 and GATA6 co-occupancy was also detected at the RE78 (Gjaltema *et al*, 2022), a locus where GATA6 is not bound in XEN cells (Fig. 5D). At RE78, *Nr5a2* KD resulted in reduced chromatin accessibility and H3K27ac, suggesting that this putative enhancer is activated through NR5A2-mediated pioneering activity. Although it remains to be tested whether RE78 directly regulates *Xist*, these data suggest that NR5A2 regulates *Xist* expression, either directly or indirectly, through its role in up-regulating GATA factor expression (Fig. 5E).

### NR5A2 co-binds to nucleosomes with KLF5 and GATA6 *in vitro*

Our genomics analysis suggest that NR5A2 co-occupies chromatin with KLF5 and GATA6 to regulate transcriptional activation at the 8-cell stage. However, bulk CUT&Tag data cannot distinguish between whether a site is simultaneously bound by factors or whether the site is occupied by one factor in some cells and another factor in other cells. To test whether KLF5 and GATA6 can principally co-bind to the same nucleosome with NR5A2, we purified recombinant full-length mouse NR5A2 as well as DNA-binding domains (DBD) of KLF5 and GATA6 (Fig. 6A). To test their DNA-binding specificities, we performed electrophoretic mobility shift analysis (EMSA). KLF5 DBD preferentially bound to the naked DNA containing its own motif, though some non-specific binding was observed (Fig. EV5A). On the other hand, GATA6 DBD bound specifically to its own motif (Fig.EV5A), confirming that the purified proteins selectively recognize their target DNA sequences.

**Figure 6.**
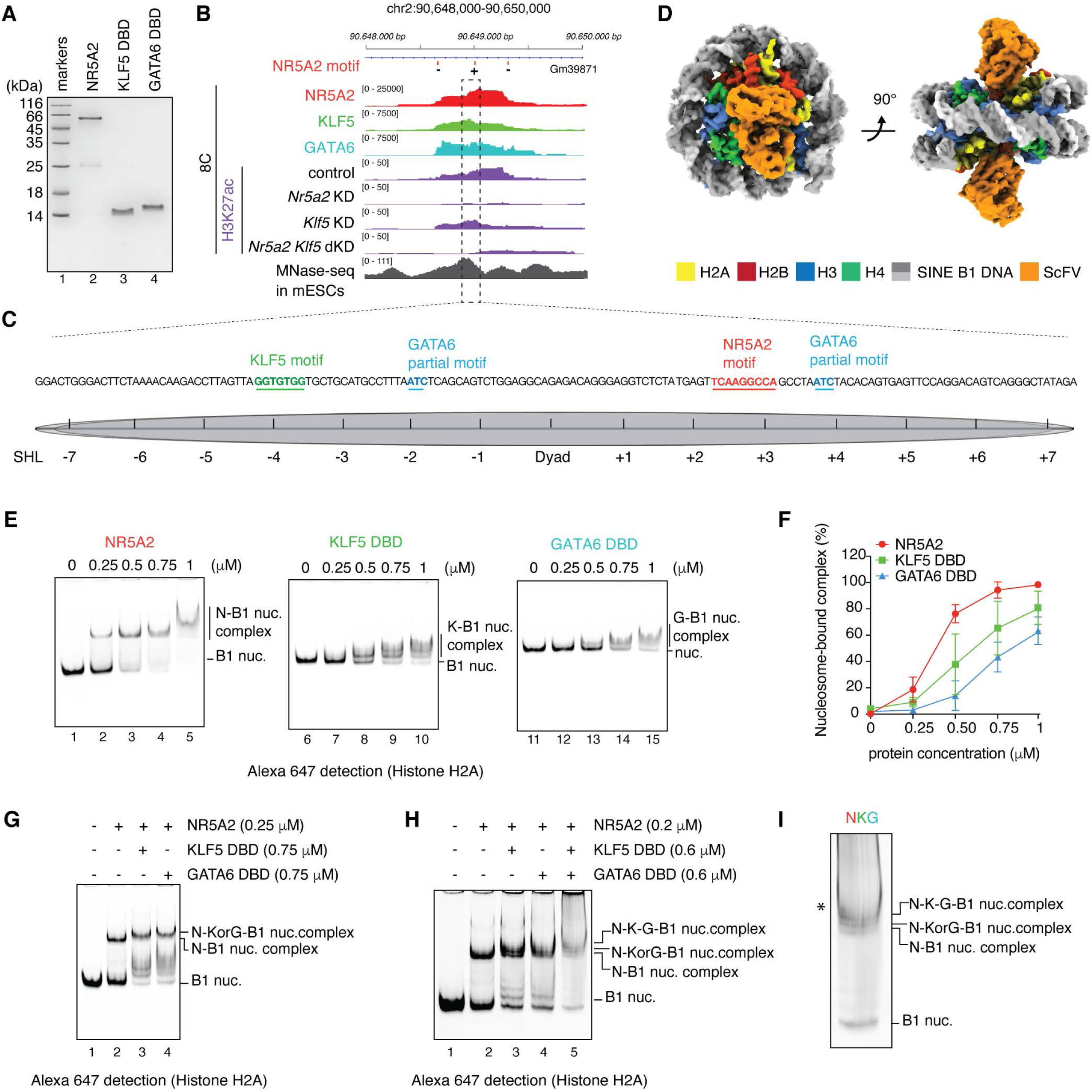
Nucleosome binding by pioneer transcription factors NR5A2, KLF5, and GATA6. (**A**) Purified mouse NR5A2 full-length (lane 2), KLF5 DBD (lane 3), and GATA6 DBD (lane 4). Lane 1 indicates molecular markers. (**B**) IGV snapshot showing co-occupancy by NR5A2, KLF5, and GATA6 along with MNase-seq profiles in mESCs. The dashed rectangle highlights the region used for mono-nucleosome reconstitution. (**C**) The sequence of 149bp DNA fragment containing SINE B1/Alu used for mono-nucleosome reconstitution. NR5A2 (red), KLF5 (green), and GATA6 (blue) motifs are highlighted. (**D**) The cryo-EM structure of B1-nucleosome bound by ScFV. (**E, F**) Representative gel images (**E**) and quantifications (**F**) of EMSA. Nucleosomes were analyzed by 5% native-PAGE and detected by Alexa Flour 647 fluorescence. Data are shown as mean ± s.d for three independent experiments. (**G**) Representative EMSA gel showing TFs co-binding on the B1-nucleosome. The bands correspond to NR5A2-nulceosome complex, NR5A2-KLF5 DBD-nucleosome complex. NR5A2-GATA6 DBD-nucleosome complex. Nucleosomes were analyzed by 6% native-PAGE and detected by Alexa Flour 647 fluorescence. Reproducibility was confirmed with three independent experiments. (**H**) Representative EMSA gel showing NKG (NR5A2-KLF5-GATA6) co-binding on the B1-nucleosome. Nucleosomes were analyzed by 6% native-PAGE and detected by Alexa Flour 647 fluorescence. Reproducibility was confirmed with three independent experiments. (**I**) Enlarged image of lane 5 from panel H. An asterisk marks an additional band, potentially non-specific KLF5 DBD binding to the NKG-B1 nucleosome complex.

To examine direct interactions between nucleosomes and NR5A2, KLF5 and GATA6 (hereafter NKG), we selected a nucleosome-enriched region measured by MNase-seq in mESCs and targeted by NKG at the 8-cell stage (Fig. 6B). *Nr5a2* KD and *Nr5a2 Klf5* dKD resulted in reduced H3K27ac, indicating that NR5A2 contributes to establishing active chromatin at this genomic locus. The selected DNA sequence of 149 bp is a part of *SINE B1/Alu* that contains NR5A2 motif (5’-TCAAGGCCA-3’) and KLF5 motif (5’-GGTGTGG-3’), and two separated GATA6 partial motifs (5’-GAT-3’) that can be recognized by one of tandem zinc finger domains (Fig. 6C). We reconstituted the nucleosome containing selected *SINE B1* DNA, resulting in NR5A2, KLF5, and GATA6 motifs located near superhelical location (SHL) +2.5, −4, and −2/+4, respectively. Single-particle analysis using cryo-EM was performed in the presence of the single-chain variable fragment (ScFv) of a nucleosome antibody, which can stabilize the nucleosomes by binding to the acidic patch regions on histones (Zhou & Bai, 2021). We determined the nucleosome containing selected mouse endogenous DNA at 4.1 Å resolution without crosslinking (Fig. 6D and EV5B-E). The nucleosomal DNA was tightly wrapped around histones, showing that the nucleosome containing *SINE B1* (hereafter referred to as B1 nucleosome) forms a canonical nucleosome structure.

We then tested NKG binding specificity to the B1 nucleosome. We detected specific band shifts corresponding to the N, K, or G-B1 nucleosome complex, suggesting that the selected endogenous nucleosome can be targeted by these three TFs *in vitro* (Fig. 6E, F and EV5F). Competition assays revealed that NR5A2 or GATA6 DBD-B1 nucleosome complex was displaced by x10-20 excess specific competitor DNA, but not in the same amount of non-specific competitor DNA (Fig. EV5F, lanes 1-6 and 17-22). In contrast, the x20 excess of non-specific competitor DNA largely outcompeted KLF5 DBD from the nucleosome (Fig. EV5G, lane 16), while x2-10 excess of non-specific competitor DNA still allowed detectable KLF5 DBD-B1 nucleosome complex (Fig. EV5G, lanes 7-15). Thus, NR5A2, KLF5, and GATA6 directly engage with their own motifs on nucleosomal DNA.

To determine whether NR5A2 co-binds nucleosomes with either KLF5 or GATA6, KLF5 DBD or GATA DBD (0.75 μM) was incubated with the B1 nucleosome in the presence of a low amount of NR5A2 (0.25 μM). Higher-order complexes with slower migration than the NR5A2-B1 nucleosome complex were detected (Fig. 6G, lanes 2-4), suggesting that NR5A2 co-binds to the nucleosome with KLF5 or GATA6. Lastly, we tested whether NR5A2 co-binds with both KLF5 and GATA6 to the B1 nucleosome using a slightly reduced concentration of each transcription factor to limit the total amount of protein. Higher-order complexes with slower migration than the NR5A2-KLF5 and NR5A2-GATA6 complexes were detected upon addition of all three transcription factors (Fig. 6H lane 5 and I). These *in vitro* binding assays suggest that the pioneer factor NR5A2, KLF5 and GATA6 can simultaneously target nucleosomes containing their recognition motifs *in vitro*.

## Discussion

In this study, we show that NR5A2 regulates the expression of KLF and GATA family transcription factors, which in turn function as co-regulators of NR5A2 during the totipotency-to-pluripotency transition. KLF5 cooperates with NR5A2 by enhancing H3K27ac deposition at regulatory elements, while GATA6 binds to distal *Xist* enhancers in conjunction with NR5A2. Thus, NR5A2 is key factor that orchestrates gene regulatory networks involved in lineage-determining factors prior to cell fate commitments. Our findings highlight how feed-forward regulatory loops by NR5A2 ensure robust gene activation during pre-implantation development (Fig. 7A).

**Figure 7.**
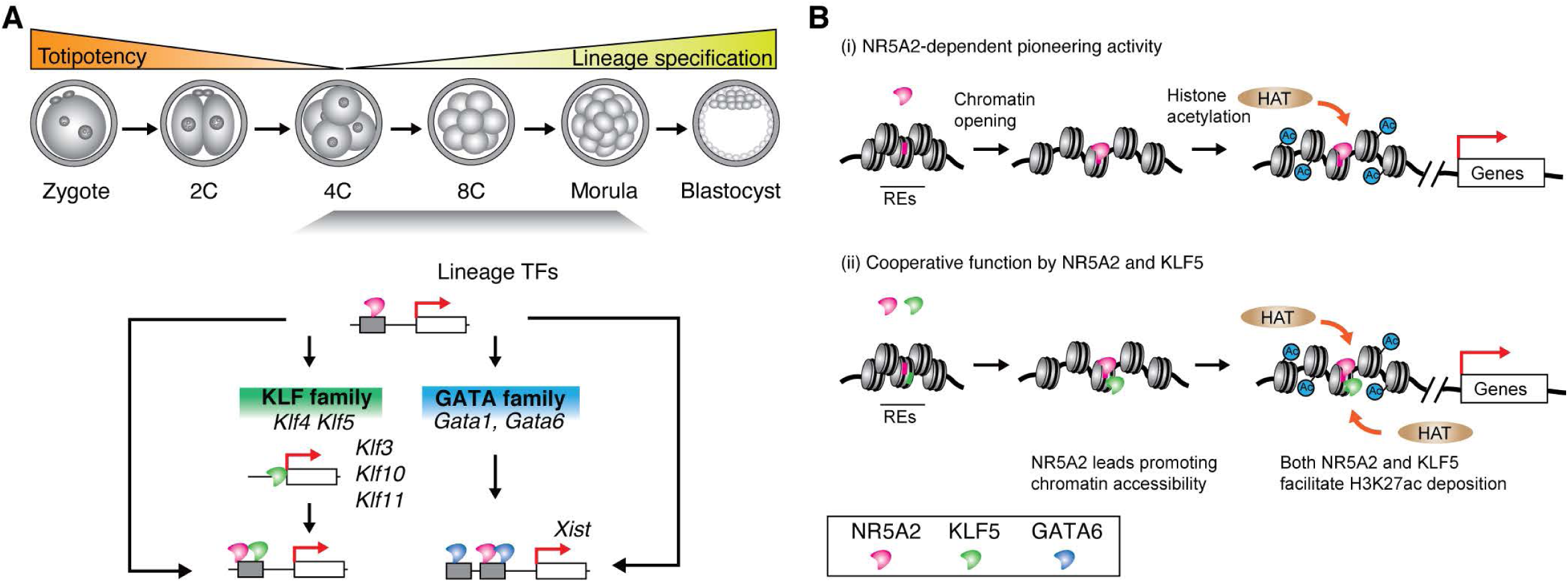
Feed-forward loop regulation by NR5A2 during the totipotency-to-pluripotency transition. (**A**) Model of feed-forward loop regulation by NR5A2. After ZGA, NR5A2 regulates lineage-determining factors including KLF and GATA families that regulate lineage commitment. KLF5 further up-regulates other KLF family: Klf3, Klf10, and Klf11. KLF5 and GATA6 co-occupy the chromatin with NR5A2 and regulate transcription. (**B**) Proposed models of pioneer factor function of NR5A2. (i) NR5A2 recognize and binds to its own motif sequence on nucleosomal DNA. NR5A2 locally opens the closed chromatin and further recruits histone acetyltransferase (HAT) such as p300 to establish active enhancers. (ii) At the NR5A2 KLF5 co-occupied regions, NR5A2 leads increasing chromatin accessibility. Following local chromatin opening, both NR5A2 and KLF5 recruit HAT to enhance histone acetylation for transcriptional activation.

Our genomics assays reveal that NR5A2 promotes chromatin accessibility and facilitates H3K27ac deposition as a pioneer transcription factor. However, the exact mechanisms by which NR5A2 leads to chromatin organization and regulates epigenetic states remains unknown. Our Cryo-EM analysis of NR5A2-nucleosome complex reveals that NR5A2 partially unwraps nucleosomal DNA at entry-exit site (Kobayashi *et al*., 2024). This structural change on the nucleosome is presumably required for local chromatin opening by evicting linker histone H1. Given that orphan nuclear receptors are associated with ATP-dependent chromatin remodelers and histone acetyltransferase p300 (Adachi *et al*, 2018; Chervova *et al*, 2024), NR5A2 may recruit these factors to disrupt the nucleosome and promote deposition of active chromatin marks at regulatory elements (Figure 7B, (i)).

In contrast, KLF5 enhances H3K27ac, but has less effect on chromatin accessibility. KLF5 is required for proper pre-implantation development and regulates lineage specification (Ema *et al*., 2008; Kinisu *et al*., 2021; Lin *et al*., 2010). In trophoblast stem cells, KLF5 maintains opening chromatin and deposits H3K27 at transcription starts sites through interaction with p300 (Dou *et al*, 2024). Despite its importance, KLF5 perturbation has relatively little effects in 8-cell embryos, possibly due to redundant function of KLF4 or other KLF family members (Kinisu *et al*., 2021). We were not able to address whether NR5A2 facilitates KLF5 chromatin binding because KLF5 expression is regulated by NR5A2. KLF5 DBD efficiently co-binds to B1 nucleosomes with NR5A2 on nucleosomes *in vitro,* supporting that NR5A2 and KLF5 independently target their own motifs in the chromatin. Hence, we propose that NR5A2 primarily opens chromatin accessibility and deposits H3K27ac in cooperation with KLF5 (Fig. 7B (ii)).

A notable finding is that the NR5A2-binding is highly dynamic and undergoes changes with each blastomere division. In addition, NR5A2 targeting to *SINE B1*/*Alu* elements gradually decreases at later developmental stages. The mechanisms by which pioneer transcription factors selectively bind their own motifs in the chromatin remains an open question. Previous studies have proposed that an increase in pioneer transcription factor concentration correlates with chromatin opening and binding at new genomic regions (Blassberg *et al*, 2022; Gibson *et al*, 2024). Consistent with this, the number of NR5A2 peaks gradually increases concomitantly with higher RNA expression level at the 8-cell stage, suggesting that local high concentration of NR5A2 may increase the chance of chromatin binding events and keep chromatin opening. Another possibility involves the regulation of *SINE B1*/*Alu* accessibility. The shutdown of SINEB1/Alu may modulate NR5A2 binding distribution. It has been reported that KRAB (Krüppel associated box) zinc finger protein ZFP266 binds to *SINE B1*/*Alu* and impedes chromatin opening by pluripotent factors (Kaemena *et al*, 2023). Though RNA expression of *Zfp266* in mouse embryos is low (Fig. EV2G), KRAB zinc finger protein may play a specific role in repressing *SINE B1*/*Alu* element during pre-implantation development.

Recently, a model has been proposed in which motif grammar on nucleosome fibers, termed as signpost elements, directs TF combinatorial binding during cell reprogramming (O’Dwyer *et al*, 2025). It would be interesting to decode motif grammar governing the combination of nuclear receptors, KLF and GATA factors during development. Taken together, our study revealed the dynamics of chromatin binding profiles of TFs during the totipotency-to-pluripotency transition and provided fundamental insights into how transcription factors ensure robust gene activation through feed-forward loops.

## Acknowledgements

We thank R. Hornberger and C. Kobayashi for technical assistance. We thank D. Bollschweiler and T. Schäfer at the cryo-EM facility (RRID:SCR_025744) for assistance in cryo-EM data collection, R. H. Kim, A. Casper, and R. Gautsch for sequencing at the NGS facility (RRID:SCR_025746), and animal facility. K.T. is an Honorary Professor at the Department of Biology, Ludwig-Maximilians-University, Munich, Germany.

## Author contributions

WK performed microinjection with micromanipulator, except for Gata6 KD, which was performed by AM. WK and EA performed genomic assays. SR performed the bioinformatic analysis. WK and EA performed biochemical experiments. WK performed cryo-EM analysis. KT conceived the project and supervised the work. WK, SR, and KT planned the project, designed the experiments, and wrote the manuscript. All authors discussed the results and commented on the manuscript.

## Data and materials availability

The raw sequencing data and processed data of this paper have been deposited in the Gene Expression Omnibus (GEO) database under the accession number: GSE289580. The dataset of NR5A2 ChIP-seq were obtained from GSE92412. MNase-seq data in mESCs was obtained from GSE82127. Datasets from the following accessions were used as inputs for ChromHMM: GSE72784, GSE73952, GSE71434, and GSE98149. Requests for plasmids generated in this study should be directed to the corresponding author.

**Figure EV1. Mapping of NR5A2 chromatin binding and chromatin states**

**(A)** IGV snapshot showing ATAC-seq and H3K27ac CUT&Tag signals across developmental stages: 2-cell (blue), 4-cell (purple), 8-cell (red), and morula (green). Publicly available of H3K4me3 ChIP-seq data (Liu *et al*, 2016) is shown. (**B, C, D**) Correlation matrix (Pearson correlation) of NR5A2 CUT&Tag (**B**), ATAC-seq (**C**), and H3K27ac CUT&Tag (**D**) comparing between replicates and cell stages. (**E**) Chromatin state annotation of by ChromHMM. (**F**) *De novo* motifs of NR5A2 peaks in different clusters. P-values and percentage of motif presence in peaks are shown.

**Figure EV2. Testing of siRNA-mediated knockdown**

**(A)** Representative images of immunostaining analysis showing GATA6 (grey) and DAPI (cyan) in 8-cell embryos. Both signals are shown in mid-section images. The numbers of embryos examined (n) from one experiment are indicated. (**B**) Abundance of co-injection marker *H2B-EGFP*, *Nr5a2*, and *Klf5* transcripts in control (blue), *Nr5a2* KD (pink), *Klf5* KD (yellow), and *Nr5a2 Klf5* dKD (purple) 8-cell embryos. (**C**) Principal component analysis of RNA-seq data in control and KD 8-cell embryos. (**D**) Pearson correlation analysis of ATAC-seq and H3K27ac CUT&Tag in control and KD 8-cell embryos. (**E**) Principal component analysis of H3K27ac CU&Tag and ATAC-seq data in control and KD 8-cell embryos. (**F**) IGV snapshot highlighting the representative early ICM and TE genes regulated by NR5A2. ATAC-seq and H3K27ac CUT&Tag are shown as merged profiles. (**G**) Expression level of KRAB (Krüppel associated box) zinc finger protein ZFP266 (see Discussion).

**Figure EV3. KLF5 regulates other KLF family members at the 8-cell stage**

**(A)** Heatmaps showing z-normalized value (*Klf5* KD/control) in gene expression. KLF family members are highlighted in red, (**B**) Abundance of *Klf3*, *Klf4, Klf10,* and *Klf11* transcripts in control (blue), *Nr5a2* KD (pink), *Klf5* KD (yellow), and *Nr5a2 Klf5* dKD (purple) 8-cell embryos. (**C**) Bar charts showing number of nucleotides in each gene matched with *Klf5* siRNA sequence. (**D**) IGV snapshot showing KLF5 binding on transcription start sites of Klf3, Klf10, and Klf11 genes.

**Figure EV4. Non-coding RNA *Xist* is regulated by NR5A2**

**(A)** Prediction of sex type by detectable Y chromosome gene expression. The clusters of red and turquoise corresponds to predicted female and male, respectively. (**B**) Box plots showing *Xist* and XCI-related genes in control, maternal-zygotic (mz) *Nr5a2 Esrrb* KO, and mz *Nr5a2* KO 8-cell embryos.

**Figure EV5. Cryo-EM data processing and motif specificity of NKG.**

**(A)** DNA-binding specificity of KLF5 DBD (left) and GATA6 DBD (right). Three independent experiments were performed, and the reproducibility was confirmed. (**B**) Flowchart of the dataset obtained by Glacios. Cryo-EM map of B1 nucleosome determined by RELION. (**C**) Local resolution of map of B1 nucleosome structure. (**D**) Fourier Shell Correlation (FSC) curves of map. The resolution of the final 3D map was 4.09 Å, as estimated by the gold standard at FSC = 0.143. (**E**) Angular distribution plot of particles employed to reconstruct maps. (**F**) SYBR gold staining of the gel shown in Figure 6E. (**G**) Competition assay testing binding specificity. NR5A2, KLF5 DBD, or GATA6 DBD was incubated with B1 nucleosome (50 nM) in the presence of their specific competitor DNA (“s” lanes) or non-specific DNA (“ns” lanes) with indicated amounts. Three independent experiments were performed, and the reproducibility was confirmed.

**Table S1. NR5A2 peaks from 2C to morula stages.**

Information of NR5A2 peaks in 2C, 4C, 8C, and morula. The mouse genomic loci at NR5A2 peaks are shown.

**Table S2. ChromHMM input lists.**

Public data used for ChromHMM analysis. Accession numbers and cell type are shown.

**Table S3. The list of differentially expressed genes.**

Results of DESeq2 analysis comparing control and knockdown embryos.

## Material and methods

### Animals

All animals housed at MPIB were sacrificed prior to the removal of organs in accordance with the European Commission Recommendations for the euthanasia of experimental animals (Part 1 and Part 2). Breeding and housing as well as the euthanasia of the animals are fully compliant with all German (e.g. German Animal Welfare Act) and EU (e.g. Directive 2010/63/EU) applicable laws and regulations concerning care and use of laboratory animals. Mice were kept at a daily cycle of 14-h light and 10-h dark with access to food ad libitum. Mice were bred in the MPIB animal facility.

### Collection of mouse embryos

We used B6CBAF1 frozen zygotes (Janvier-lab) for genome-wide mapping of NR5A2, KLF5 GATA6, H3K27ac, and ATAC-seq. Zygotes were thawed according to a manual of kits of frozen zygotes and cultured in potassium-supplemented simplex-optimized medium (KSOM) with Lumox dish (Sarstedt). Two-cell, four-cell, eight-cell, and morula were collected after thawing at 22-24 hours (34-36 hpf), 29-31 hours (41-43 hpf), 44-46 hours (56-58 hpf), and 59-60 hours (71-72 hpf), respectively. For KD experiments followed by RNA-seq, H3K27ac CUT&Tag, and ATAC-seq, zygotes were collected from superovulated B6129F1 female mice (3-5 weeks) mated with B6CBAF1male mice. To induce superovulation, females were injected with PMSG (5IU) followed by hCG injection 46-48 hours later. Zygotes were obtained by opening the oviduct after 19 hours post hCG injection.

### Immunofluorescence

Zona pellucida was removed by treatment with acidic Tyrode. Embryos were fixed in 4% formaldehyde (company) for 30 min and then permeabilized with 0.5% Triton-X in PBS. Blocking was carried out in 1% BSA (sigma, A8577) in PBTX (0.1% Triton-X in PBS) for 1 hour at room temperature. Cells were incubated with primary antibodies (NR5A2: 1:100, R&D Systems, PP-H2325-00, KLF5; 1:100, Proteintech, 21017-1-AP, GATA6: 1:500, R&D Systems, AF1700) for 1h at room temperature on nutator. Cells were washed through blocking solution three times with PBTX and were incubated with the secondary antibody (1:500, Thermo Fisher, goat anti-mouse Alexa Flour 488 or 647, goat anti-rabbit Alexa Flour 568, Donkey anti-goat Alexa Flour 594) for 30 min at room temperature. After three times washing with PBTX, cells were incubated with Vectashield + DAPI and mounted. Images were acquired by using a confocal microscope (Leica, STELLARIS 5). All images were analyzed with Fiji and arivis Pro/Vision4D. The signal intensity within nuclei of control samples was determined and the cytoplasmic signal was subtracted as background. The average signal intensity of the control samples was set as 1.0.

### Microinjection of zygotes for siRNA knockdown

For siRNA knockdown, isolated zygotes (8-9 hpf) were microinjected with siRNAs against targets *Nr5a2* (5 µM), *Klf5* (10 µM) or control (15 µM). As a control of successful injection 150 ng/µl H2B-EGFP was co-injected. The siRNA targeting control and *Nr5a2* (Lai *et al*., 2023) were ordered as *Silencer* select from Life technologies. Predesigned siRNAs were obtained from following vendors: Klf5 (#160900, Silencer pre-designed siRNA, Life technologies); To test *Gata6* KD efficiency, 100 μM of siRNA targeting *Gata6* (#158651, Silencer pre-designed siRNA, Life technologies) was injected into zygotes. The injected embryos were cultured in KSOM using lumox dishes (35 mm, Sarstedt) under 5% O_2_, 5% CO_2_, 90% N_2_.

### Embryonic development assay

Embryos were cultured in 30 µl microdrops of KSOM using lumox dishes (35 mm, Sarstedt) under 5% O_2_, 5% CO_2_, 90% N_2_. Embryonic development was monitored once every 24 hours using a stereomicroscope with a camera (Flexacam C3, Leica Microsystems).

### CUT&Tag

CUT&Tag was performed as described previously (Gassler *et al*., 2022). Embryos with intact zona pellucida were incubated with ice-cold extraction buffer (25 mM HEPES-NaOH (pH 7.4), 50 mM NaCl, 3 mM MgCl_2_, 300 mM sucrose and 0.5% Triton X-100) on ice for 6-7 min and were washed three times through with ice-cold extraction buffer without TritonX-100. Pre-extracted embryos were immediately lightly fixed by Dulbecco’s phosphate-buffered saline (DPBS) with 0.1% formaldehyde for 2 min at room temperature. Embryos were incubated in antibody buffer (20 mM HEPES-NaOH (pH7.5), 150 mM NaCl, 0.5 mM Spermidine, 0.1% BSA, 2 mM EDTA and 1x Protease inhibitor cocktail (Roche) with primary antibody (1:100 dilution, NR5A2 (R&D Systems, PP-H2325-00), KLF5 (Proteintech, 21017-1-AP), GATA6 (R&D Systems, AF1700), H3K27ac (Active motif, 39133)) overnight at 4°C on nutator. Embryos were washed three times through antibody buffer and further incubated with secondary antibody (1:100 dilution, Guinea pig anti-rabbit H&L antibody (ABIN101961, Antibodies-Online) for anti-rabbit primary antibody, rabbit anti-mouse IgG H&L (ab46540, abcam) for anti-mouse primary antibody, and Rabbit anti-goat IgG H&L (ab6697, abcam) for anti-goat primary antibody) for 1.5 hours at room temperature. Cells were washed three times through Wash buffer (20 mM HEPES-NaOH (pH7.5), 150 mM NaCl, 0.5 mM Spermidine and 1x Protease inhibitor cocktail (Roche)) and further incubated with homemade pA-Tn5 or pAG-Tn5 adaptor complex (1:250 dilution) for 1.5 hours at room temperature. Embryos were washed three times through Wash-300 buffer (20 mM HEPES-NaOH (pH7.5), 300 mM NaCl, 0.5 mM Spermidine and 1x Protease inhibitor cocktail (Roche)). Embryos were transferred into 1.5 ml of DNA LoBind Tube (Eppendorf) with 200 µl of Tagmentation buffer (10 mM MgCl_2_ in Wash-300) and incubated at 37°C for 1 hour. The samples were then deproteinized and reverse-crosslinked. The DNA was purified by phenol-chloroform and followed by ethanol precipitation. The DNA was amplified 50 µl of reaction with 1×Q5 High-Fidelity Master mix (NEB) and 0.25 µM barcoded primers using the following PCR program: 72 °C for 5 min; 98°C for 30 sec; 15 cycles of 98°C for 10 sec and 63°C for 10 sec; final extension at 72 °C for 1 min and hold at 4°C. After PCR reaction, libraries were purified by 1.1×AMPure XP magnetic beads (Beckman Coulter) and were eluted in 20 µl of RNase-free water. Purified DNA libraries were sequenced on a NextSeq 500 (Illumina), Novaseq 6000 (Illumina), or AVITI (Element Biosciences) with paired-end reads.

### Omni ATAC-seq

Omni ATAC-seq was performed as described previously (Gassler *et al*., 2022). Zona pellucida was removed by treatment with acidic Tyrode. Embryos were transferred into 1.5 ml of DNA LoBind Tube (Eppendorf) with 200 μl of PBS and were spun at 500×g at 4°C for 5 min to remove supernatant. 50 μl of ATAC-Resuspension buffer (RSB) (10 mM Tris-HCl (pH 7.4), 10 mM NaCl, 3 mM MgCl_2_) containing 0.1% NP-40, 0.1% Tween-20 and 0.01% Digitonin was added into tube and incubated for 3 min on ice. After incubation, 200 μl of ATAC-RSB containing 0.1% Tween-20 was further added into the tube to wash out lysis. Nuclei were spun at 500×g at 4°C for 10 min to remove supernatant. Nuclei were incubated with 50 μl of transposition mixture (10 mM Tris-HCl (pH 7.6), 5 mM MgCl_2_, 10% Dimethyl formamide, 33% PBS, 0.1 % Tween-20, 0.01% Digitonin and 95 nM Tn5-adaptor complex) at 37°C for 30 min with 1000 rpm mixing. The samples were then deproteinized and further purified by phenol-chloroform and followed by ethanol precipitation. The DNA was amplified 50 µl of reaction with 1×NEBNext HF 2×PCR Master mix (NEB) and 0.25 µM barcoded primers using the following PCR program: 72 °C for 5 min; 98°C for 30 sec; 15 cycles of 98°C for 10 sec, 63°C for 30 sec and 72 °C for 1 min; final extension at 72 °C for 5 min and hold at 4°C. After PCR reaction, libraries were purified by 1.1×AMPure XP magnetic beads (Beckman Coulter) and were eluted in 20 μl of RNase-free water. Purified DNA libraries were sequenced on a NextSeq 500 (Illumina), or AVITI (Element Biosciences) with paired-end reads.

### RNA-seq library preparation

The zona pellucida was removed by treatment with acidic Tyrode. 10 x 8-cell embryos were washed five times in M2 medium and then quickly washed in PBS. Embryos were lysed in lysis buffer containing RNase inhibitor and ERCC spike-in RNA on ice for 15min and then snap frozen with liquid nitrogen. cDNA was prepared using a SMART-seq v4 Ultra-Low-Input RNA kit for Sequencing (Takara, 634888). Library was prepared by a Nextera XT DNA Library Prep Kit (Illumina, FC-131-1024) according to the manufacturer’s protocol. Purified DNA libraries were sequenced on a Novaseq 6000 (Illumina) with paired-end reads.

### Protein purification

His_6_-tagged full-length mouse NR5A2 was purified as described previously (Gassler *et al*., 2022). The DNA fragment encoding *Mus musculus* KLF5 DBD (348-446) was ligated into *Bam*HI-*Not*I sites of pGEX6P-1 vector. Glutathione-S-transferase (GST)-fused KLF5 DBD was expressed in *E.Coli* BL21 (DE3) codon plus RIL (Agilent Technologies). The cells were cultivated at 30°C, and the protein expression was induced with 0.25 mM IPTG at 16°C for 16-18 h. The cells were resuspended in buffer 1 (50 mM Tris-HCl (pH 7.5), 500 mM NaCl, 1 mM EDTA, 10% glycerol, and 2 mM 2-mercaptoethanol). The cells were disrupted by sonication, and the cell debris was further removed by centrifugation. The protein bound to the Glutathione sepharose 4B beads (Cytiva) was washed by 50 column volumes (CVs) with buffer 1. GST-tag was cleaved with homemade PreScission protease on the column at 4°C overnight. The supernatant containing KLF5 DBD was collected and concentrated with an Amicon Ultra centrifugal filter unit (Millipore).

*Mus musculus* GATA6 DBD was purified as described previously with a few modifications (Takaku *et al*, 2016). The codon-optimized DNA fragment encoding *Mus musculus* GATA6 DBD (382-492) was ligated into *Nde*I-*Bam*HI sites of pET-15b vector with a N-terminal His_6_-TEV cleavage site. His_6_-tag GATA6 DBD was purified by Ni-NTA agarose beads (Qiagen). His_6_-tag was removed by homemade TEV protease, and GATA6 DBD was purified by MonoS 5/50 GL (Cytiva) column chromatography. The sample was further loaded onto Superdex200 16/60 (Cytiva), and fractions containing GATA6 DBD were stored at −70°C.

Mouse histones H2A K119C, H2B, H3.3, and H4 were expressed and purified as previously described (Kujirai *et al*, 2018). Histone H2AK119C-H2B complex and H3.3-H4 complex were reconstituted as previously described (Kobayashi *et al*., 2024). For the fluorescent labeling of H2A-H2B complex, 50 μM of H2A K119C-H2B complex were incubated with 500 μM Alexa Flour 647 C_2_ Maleimide (Invitrogen) in 20 mM Tris-HCl (pH 7.5) buffer containing 0.1 M NaCl and 1 mM TCEP at room temperature for 2h in the dark. The reaction was stopped by the addition of 150 mM of 2-mercaptoethanol, and the sample was then dialyzed against 20 mM Tris-HCl buffer containing 0.1M NaCl, 1 mM EDTA, and 5 mM 2-mercaptoethanol.

The DNA fragment encoding ScFv linker 20 (ScFv^20^) was synthesized by the polymerase chain reaction (PCR) method. The amplified DNA fragment encoding ScFv^20^ was ligated into *Nde*I-*Bam*HI site in pET15b-TEV vector. ScFv^20^ was expressed and purified as previously described (Zhou & Bai, 2021).

### DNA preparations

The endogenous DNA fragment containing SINE B1/Alu for nucleosome reconstitution were amplified by PCR and further purified by polyacrylamide gel (6%) electrophoresis using a Prep Cell apparatus (Bio-Rad). The eluted DNA was concentrated with an Amicon Ultra centrifugal filter unit (Millipore).

### Preparation of the nucleosomes containing mouse endogenous DNA

The nucleosomes were reconstituted by salt dialysis method. Briefly, the DNA, Alexa 647-labeled histone H2A-H2B complex, and H3.3-H4 complex were mixed in 1:4:3.6 molar ratio in 2M KCl high salt buffer. After salt dialysis, the reconstituted nucleosomes were further purified by polyacrylamide gel (6%) electrophoresis using a Prep Cell apparatus (Bio-Rad). The nucleosomes were concentrated with an Amicon Ultra centrifugal filter unit (Millipore).

### Electrophoretic mobility shift assay (EMSA)

For EMSA with short oligo DNA (Chen *et al*, 2012; Gassler *et al*., 2022), 10% non-denaturing polyacrylamide gels (0.5xTBE) were pre-run at 100V for 30 min in the cold room. The DNA (50 nΜ) were incubated with 0-1 μM KLF5 DBD or GATA6 DBD at room temperature for 30 min in a reaction buffer (20 mM Tris-HCl (pH7.5), 120 mM NaCl, 1 mM MgCl_2_, 10 μΜ ZnCl_2_, 1 mM DTT, 100 μg/ml BSA). After the incubation, the samples were loaded onto gels, and electrophoresis was performed at 100V for 100 min in the cold room. The gels were stained by SYBR Gold (Invitrogen) and were imaged using ChemiDoc MP imaging system (Bio-Rad).

For EMSA with nucleosome, 5-6% non-denaturing polyacrylamide gels (0.5xTBE) were pre-run at 100V for 30 min in the cold room. The nucleosomes (50 nΜ) were incubated with 0-1 μM NR5A2, KLF5 DBD, or GATA6 DBD at room temperature for 30 min in a reaction buffer (20 mM Tris-HCl (pH7.5), 120 mM NaCl, 1 mM MgCl_2_, 10 μΜ ZnCl_2_, 1 mM DTT, 100 μg/ml BSA). For competition assay, the non-labeled naked DNA containing non-specific or specific sequence (0.1-1 μM) was added with the nucleosome. After the incubation, the samples were loaded onto gels and electrophoresis was performed at 100V for 80-120 min in the cold room. The gels were imaged by detecting Alexa 647 fluorescence and SYBR Gold (Invitrogen) using ChemiDoc MP imaging system (Bio-Rad).

### Cryo-EM specimen preparation and data acquisition

ScFV^20^ (1.5 μM) and the B1-nucleosome (0.5 μM) were mixed in a 3:1 molar ratio on ice for 30 min. The ScFV^20^–nucleosome complex was concentrated by Amicon Ultra centrifugal filter unit (Millipore) until 215 ng/ul (DNA). To prepare the cryo-EM specimen, the sample (4 μl) was applied to a glow-discharged holey carbon grid (Quantifoil R1.2/1.3 200-mesh Cu). The grids were blotted for 3.0 s at a blotting strength setting of 5 under 100% humidity at 4 °C and then plunged into liquid ethane and cooled by liquid nitrogen using a Vitrobot Mark IV (Thermo Fisher). Data acquisition for the ScFV^20^–B1 nucleosome complex was conducted using a 200 kV Glacios (Thermo Fisher Scientific) equipped with a GIF Quantum 967 energy filter and a K3 direct electron detector (Gatan) running in correlated double sampling mode at a magnification factor of 36kx, equivalent to a pixel size of 1.136 Å per pixel. Automated data acquisition was performed using SerialEM software. A total of 2,489 videos were collected, with a total electron dose of 60 e/Å² fractionated over 40 frames.

### Image processing

Data processing was performed by RELION 4.0 (Kimanius *et al*, 2021). In total, 2,489 micrographs were stacked and motion-corrected using MOTIONCOR2 (Zheng *et al*, 2017) with dose weighting. The estimation of contrast transfer function (CTF) from the dose-weighted micrographs was performed using CTFFIND4 (Rohou & Grigorieff, 2015). From 2,449 micrographs, 571,540 particles were automatically picked and extracted with a binning factor of 4 (pixel size of 4.724 Å/pixel). In total, 516,292 particles were further selected by 2D classification. The ab initio model generated was used as the initial model for 3D classification. After two rounds of 3D classification, 152,279 particles were selected and re-extracted without binning for 3D refinement, followed by Bayesian polishing and CTF refinement. The refined map of the ScFV^20^–B1 nucleosome complex was sharpened with a B-factor of −120 Å^2^. The resolution of the final 3D map was 4.09 Å, as estimated by the gold standard Fourier Shell Correlation (FSC) at FSC = 0.143 (Rosenthal & Henderson, 2003). This map was subsequently post-processed through local B-factor correction by DeepEMhancer (Sanchez-Garcia *et al*, 2021). The local resolution of the map was calculated by RELION4 and visualized with UCSF ChimeraX (version 1.5) (Pettersen *et al*, 2021).

### Data analysis

#### CUT&Tag, ChIP-seq, ATAC-seq data processing

Low quality reads were trimmed out using TrimGalore (version 0.6.2) (https://github.com/FelixKrueger/TrimGalore) using parameters: --paired --quality 20 –length 20. Read mapping was done using Bowtie2 (version 2.3.5.1) (Langmead & Salzberg, 2012) using parameters: -t -q -N 1 -L 25 -X 2000 --no-mixed --no-discordant. Unmapped reads and reads with low mapping quality (Q<30) were removed. PCR duplicated reads were identified and discarded using the MarkDuplicates command from Picard (version 2.18.27) (http://broadinstitute.github.io/picard/) with parameters: VALIDATION_STRINGENCY=LENIENT. The RPKM values were calculated to represent read coverage using the bamCoverage function from deepTools (version 3.5.4) using parameters: --binSize 1 --ignoreForNormalization chrM (Ramirez *et al*, 2016). The RPKM values were further transformed using Z-score normalization for the visualization using custom R script. To access similarity and reproducibility of the libraries, correlation between libraries were calculated using multiBigwigSummary command from deepTools with parameters: -- chromosomesToSkip chrM, then heatmap of library correlation were made using plotHeatmap command from deepTools with parameters: --skipZeros –removeOutliers --corMethod pearson. Heatmap of chromatin binding were generated using computeMatrix and plotHeatmap commands from deepTools. The computeMatrix command were run with the following parameters: --binSize 50 --missingDataAsZero --skipZeros.

Reads from Omni-ATAC-seq datasets were mapped to both mouse (mm10) Drosophila (dm6, spike-in) genomes. Unmapped, low mapping quality, and PCR duplicated reads were removed as described previously. Note that normalization by spike-in reads weren’t performed due to extremely low mappable spike-in reads in these samples.

For analysis of ATAC-seq and H3K27ac CUT&Tag of Nr5a2 and Klf5 knockdown samples, the log2-fold-change of chromatin coverage signals between the siRNA knockdown and control condition were calculated using bigwigCompare command from deepTools using the following parameters: --binSize 100 --skipZeroOverZero --operation log2.

The R package DiffBind (Ross-Innes *et al*, 2012; Stark & Brown, 2011) were used to generate consensus peaks, and count reads. Normalized reads counts were extracted and used for building principal component analysis in a custom R script.

#### RNA-seq data processing

Adaptor sequences were trimmed from the reads using TrimGalore (version 0.6.2) (https://github.com/FelixKrueger/TrimGalore) using parameters: --paired --quality 10 –length 10. Reads were then mapped to the custom reference transcriptome based on GRCm30 (version 102) with addition of H2B-EGFP and ERCC spike-in reference using STAR with default setting (Dobin *et al*, 2013). Expression for individual gene were quantify using featureCounts (version 2.0.1) (Liao *et al*, 2014) with the additional parameters: -t exon -g gene_id -p -O --fraction. To identify differential expressed genes (DEGs), expression data were imported to R. Genes with low counts across all samples were identified and removed using HTSFilter package (Rau *et al*, 2013). DEGs were identified using DESeq2 package (Love *et al*, 2014). Only genes with p-value cut-off of 0.05 and have at least 2-fold change compared with the control condition will further considered as DEGs.

#### Peak calling and motif analysis

Peak calling for CUT&Tag, ATAC-seq, and ChIP-seq datasets were done using MACS2’s callpeak function (version 2.2.9) with default parameters (Zhang *et al*, 2008). The identification of de novo motif analysis for transcription factor CUT&Tag peaks were done using findMotifsGenome command from HOMER (version 4.10) (Heinz *et al*, 2010) with the ‘-size given’ parameter for most of the datasets, and with ‘-size 500’ parameter for cluster analysis of Nr5a2 peaks in morula stage. The peaks lists are shown in Table S1.

#### Annotation of chromatin states

To infer chromatin state based on epigenetic status, ATAC-seq and ChIP-seq datasets of H3K4me3, H3K27ac, H3K27me3, and H3K9me3 from 2-cell embryo to mESC were collected as specified in Table S3 (Dahl *et al*, 2016; Liu *et al*., 2016; Wang *et al*, 2018; Zhang *et al*, 2016). The ChromHMM (version 1.23)(Ernst & Kellis, 2012, 2017) was used to categorize chromatin states based on these signals. The data were first binarized using BinarizeBam command with the default parameters. Then the LearnModel command was used to infer three to fifteen chromatin states based on the same input data. The results with eleven states were arbitrary chosen for further analysis. The assignment of chromatin state to active (promoter or enhancer), repressive, or open chromatin states were done based on emission signal as shown in Fig EV1.

