## Supplementary figures and images for "Feed-forward loops by NR5A2 ensure robust gene activation during pre-implantation development"

### EV1

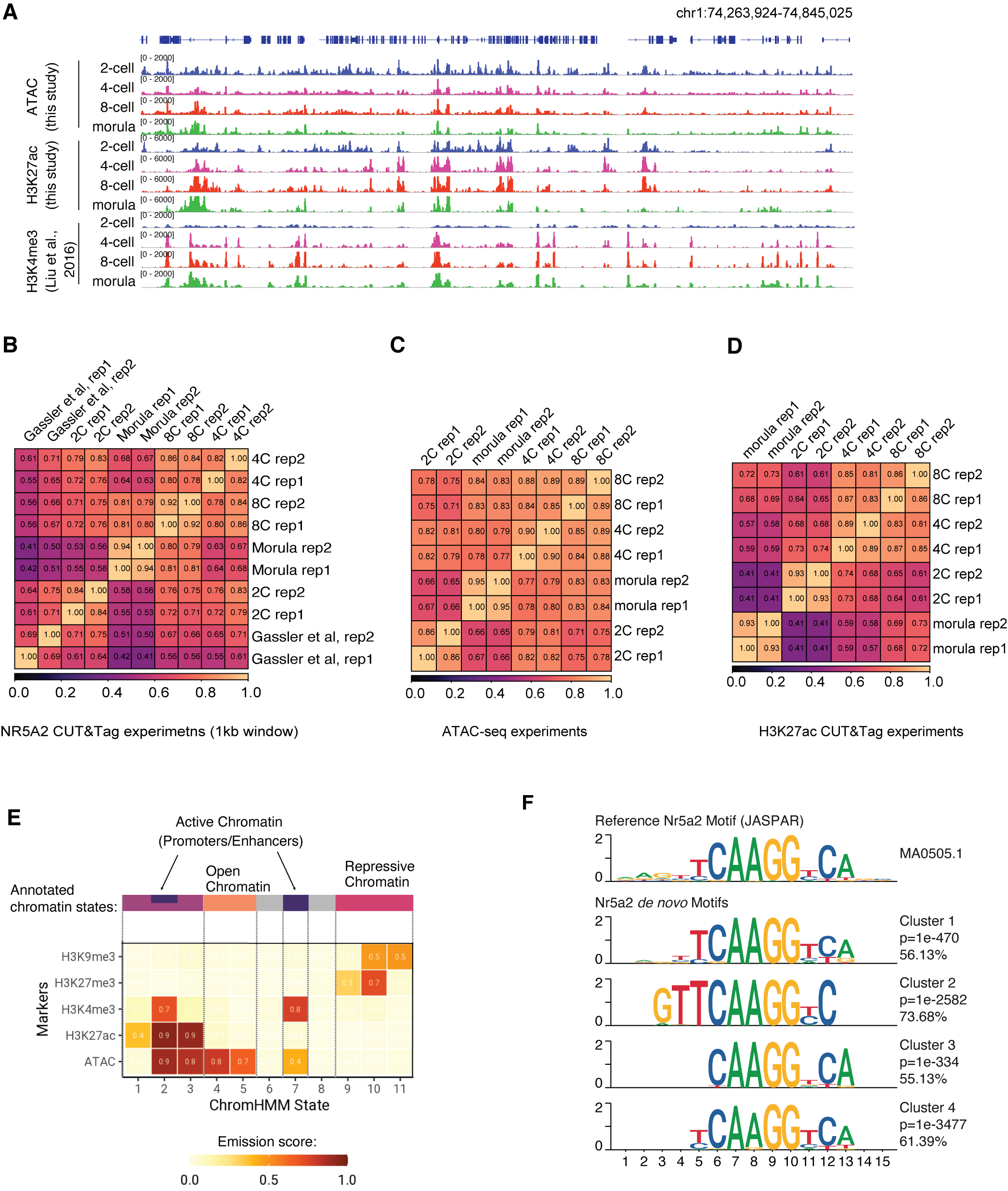

### EV2

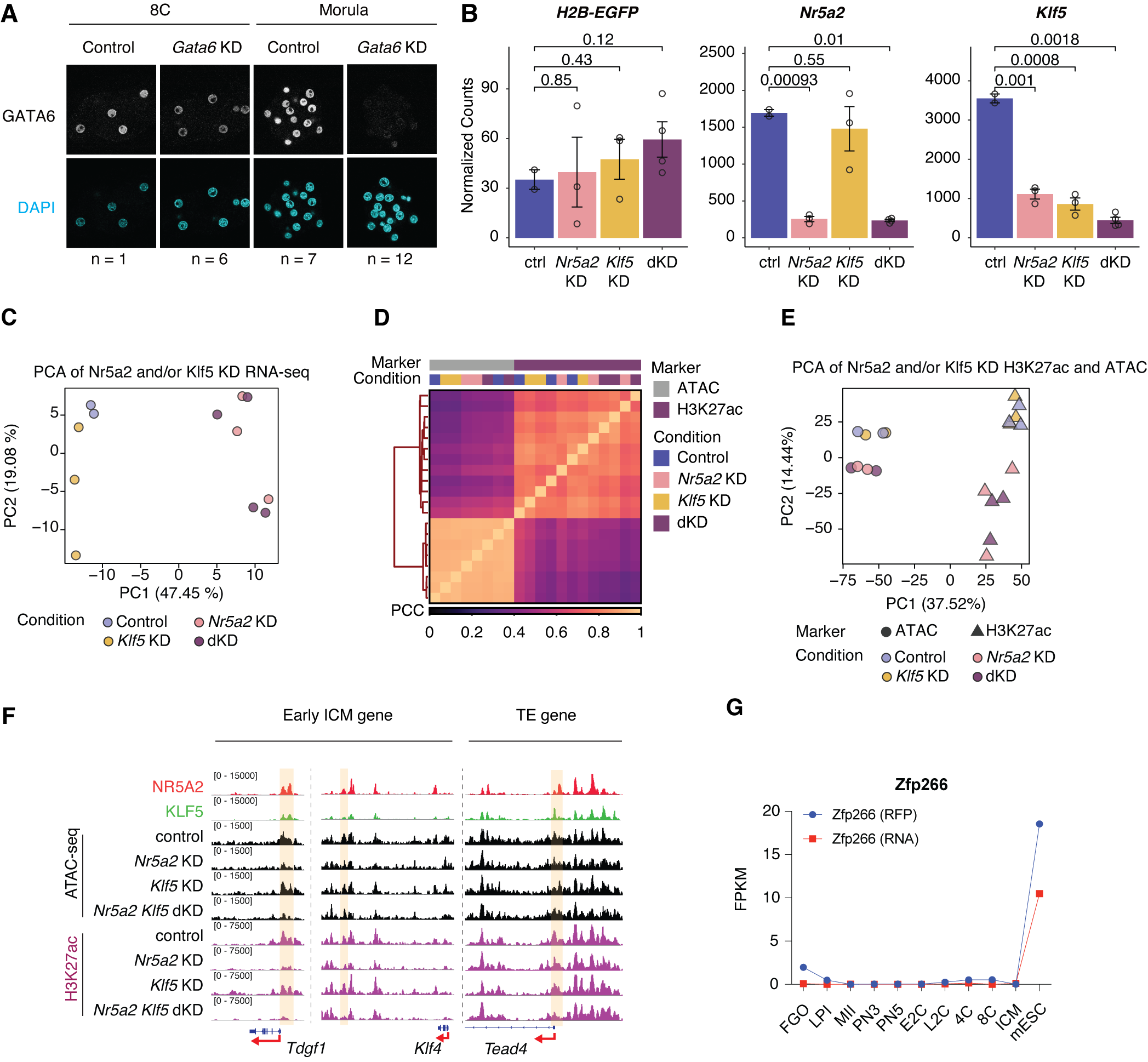

### EV3

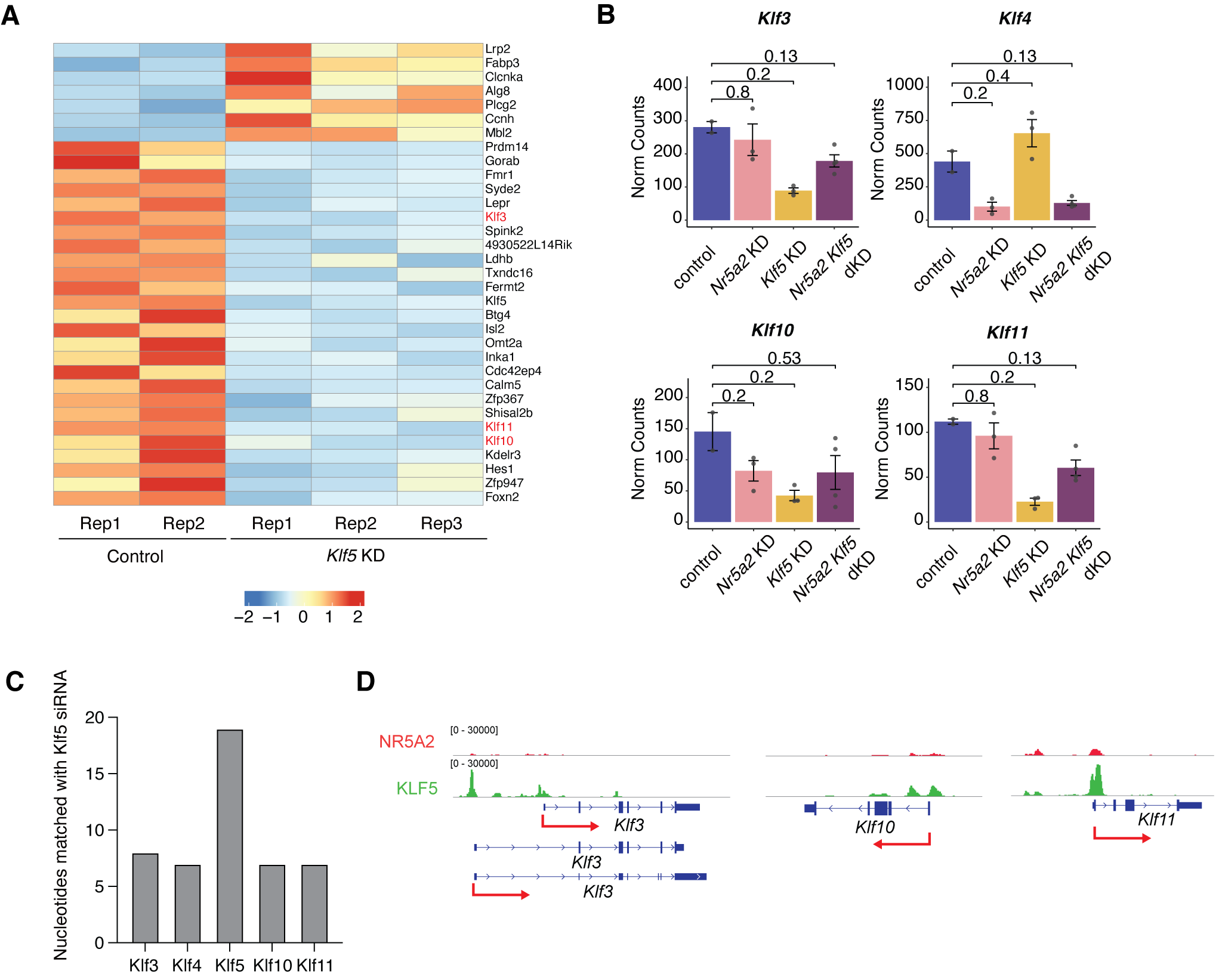

### EV4

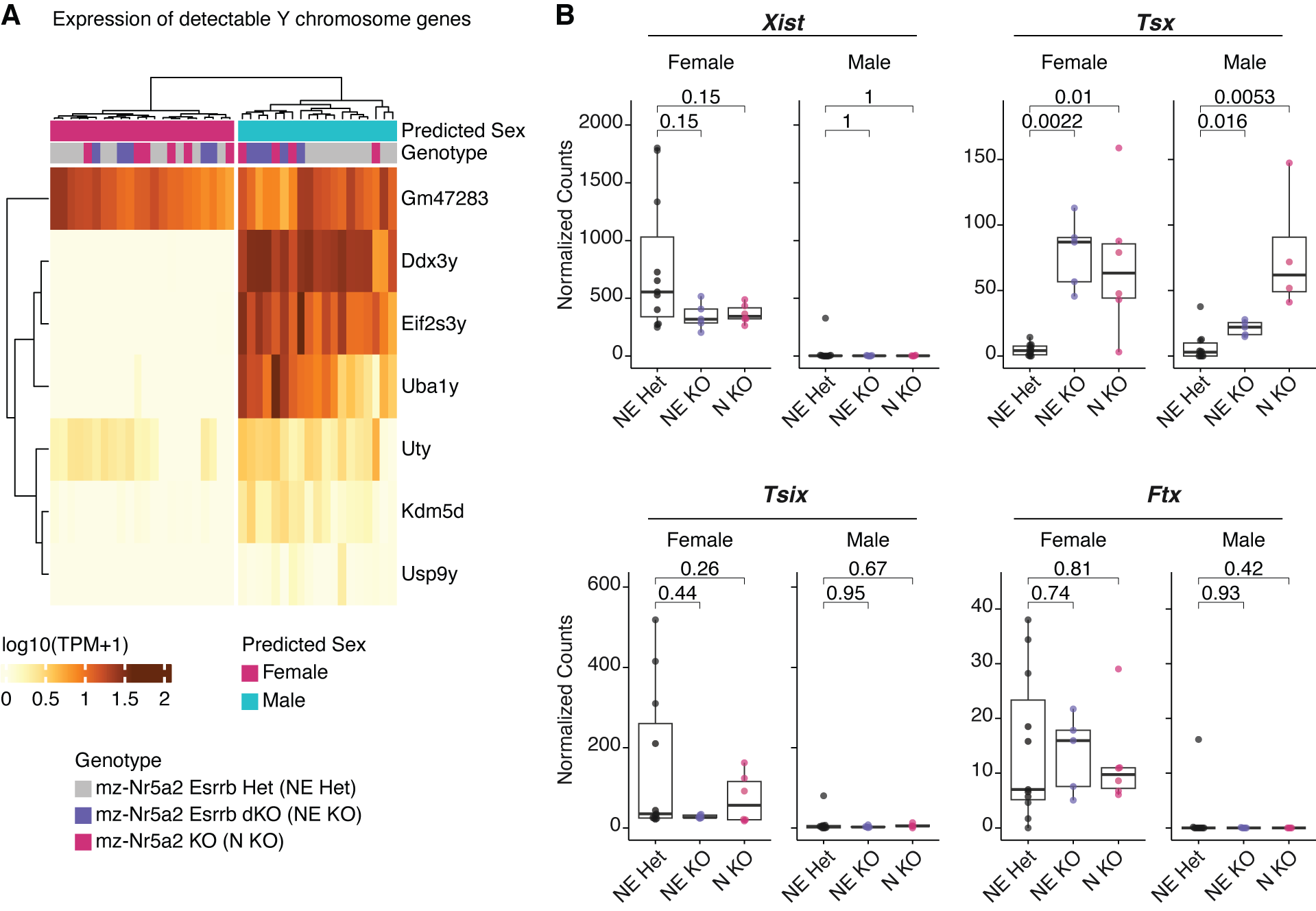

### EV5

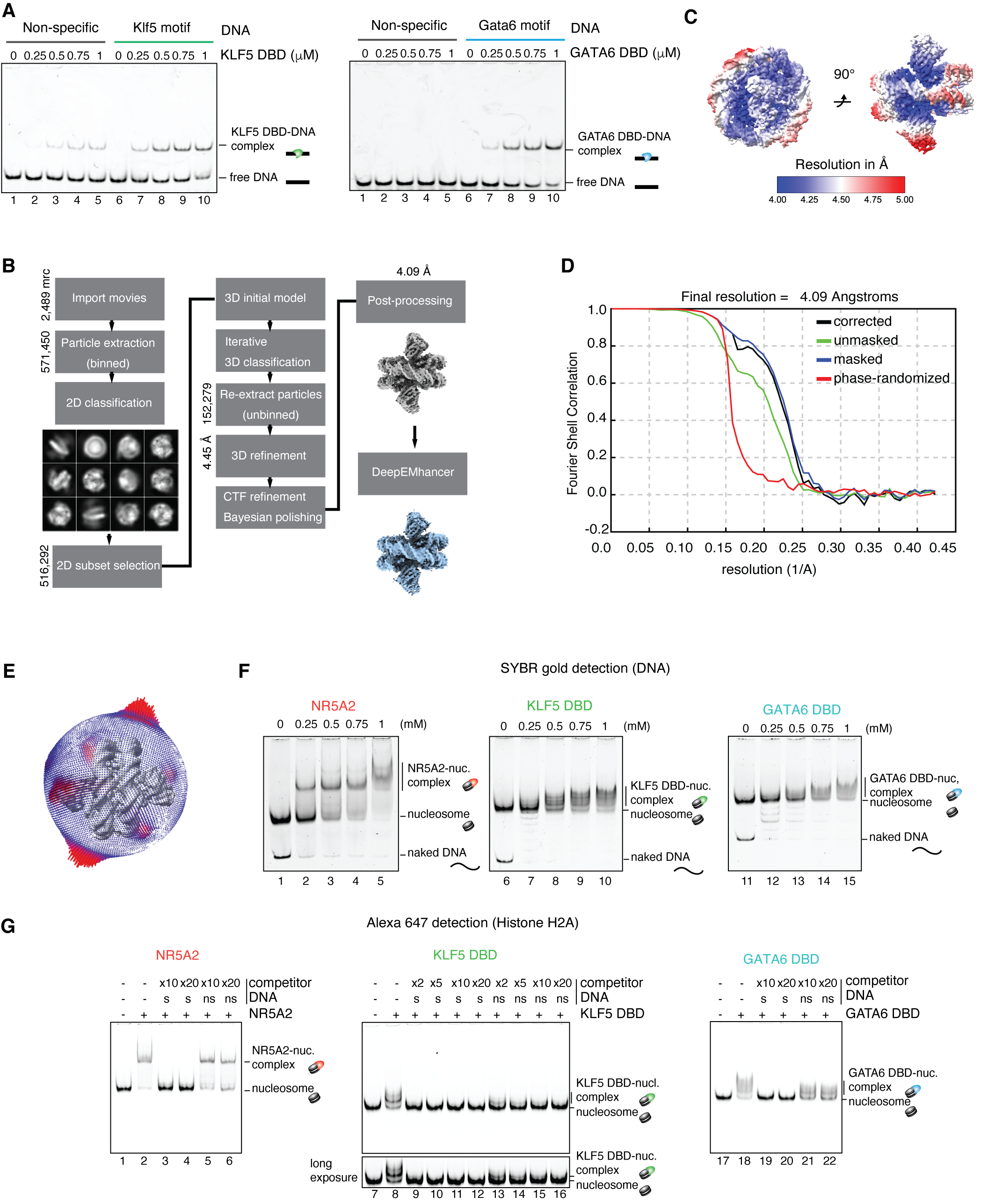
